# Substrate utilization and cross-feeding synergistically determine microbiome resistance to pathogen invasion

**DOI:** 10.1101/2024.11.04.621791

**Authors:** Xinrun Yang, Tianjie Yang, Ziru Zhang, Yaozhong Zhang, Xinlan Mei, Yang Gao, Ningqi Wang, Gaofei Jiang, Yangchun Xu, Qirong Shen, Marnix H. Medema, Zhong Wei, Alexandre Jousset

**Affiliations:** Jiangsu Provincial Key Lab for Solid Organic Waste Utilization, Key Lab of Organic-based Fertilizers of China, Jiangsu Collaborative Innovation Center for Solid Organic Wastes, Educational Ministry Engineering Center of Resource-saving fertilizers, Nanjing Agricultural University, Nanjing 210095, China; Bioinformatics Group, Wageningen University, Droevendaalsesteeg 1 Radix West, 6708PB Wageningen, The Netherlands

**Author notes:** These authors contributed equally to this work (Xinrun Yang, Tianjie Yang). **Correspondence** (Tianjie Yang), (Zhong Wei).

## Abstract

Understanding how microbiomes resist pathogen invasion remains a key challenge in natural ecosystems. Here, we combined genome-scale metabolic models with synthetic community experiments to unravel the mechanisms driving pathogen suppression. We developed curated genome-scale models for each strain, incorporating 48 common resource utilization profiles to fully capture their metabolic capacities. Trophic interactions inferred from models accurately predicted pathogen invasion outcomes, achieving an F1 score of 96% across 620 invasion tests involving diverse microbial communities and nutrient environments. Importantly, considering both substrate and metabolite features provided a more holistic understanding of pathogen suppression. In particular, cross-feeding metabolites within the native community emerged as crucial yet often overlooked predictors of community resistance, disproportionally favoring native species over invaders. This study lays the foundation for designing disease-resistant microbiomes, with broad implications for mitigating pathogen exposure in diverse environments.

## Introduction

The host-associated microbiome functions as a critical defense line against invading pathogens^1,2^. While it is well-established that high microbiome diversity limits pathogen invasion^3–6^, the underlying mechanisms remain unclear. Increasing evidence suggests that resource utilization patterns play a pivotal role in determining pathogen invasion^7,8^. Specifically, substrate competition—between the resident microbial community and the pathogen, as well as within the resident community itself—has emerged as a key factor in driving community resistance to pathogen invasion^7,9^. Despite the important role of substrate competition, the importance of microbial metabolites in pathogen suppression has largely been overlooked. Cross-feeding metabolites, defined as compounds secreted by one strain and utilized by another^10^, are key determinants of microbial interactions that may influence pathogen suppression^11,12^. Thus, understanding the dual role of substrate competition and metabolite exchange is critical to fully appreciating how microbiomes protect their host from pathogen invasion.

Predicting trophic interactions in multispecies communities has long been limited by the experimental challenges involved. *In vitro* experiments designed to examine substrate and metabolite features are already labor-intensive under controlled conditions and become increasingly impractical in complex natural settings^13^, where species composition and environmental conditions are highly variable^14,15^. Genome-scale metabolic models offer a promising solution to these challenges^16^. Using genome data, standardized pipelines can now rapidly generate metabolic models, enabling *in silico* predictions of resource utilization, metabolite secretion, and growth rates^17,18^. These models have the potential to predict metabolic interactions within community contexts, linking community composition and functional traits with susceptibility to pathogen invasion across diverse nutritional environments^19,20^.

In this study, we used defined plant-associated microbial communities to assess the predictive power of genome-scale models in explaining patterns of pathogen invasion. The plant root environment contains a host-specific set of plant-derived labile carbon compounds, which fuels the development of a host-specific microbiome^21^. These nutrients are accessible to both commensals and pathogens, and imbalances in resource utilization can promote disease^2,22^. While previous studies have primarily focused on substrate availability as a driver of pathogen invasion^7^, we hypothesize that considering both substrate and metabolite features offers a more comprehensive mechanism for pathogen suppression, with microbial cross-feeding playing an equally important but less understood role in contributing to community resistance to pathogen invasion (Fig. 1A). To test this hypothesis, we constructed resident communities of plant-associated bacteria and challenged them with invasion by the plant pathogen *Ralstonia solanacearum* (Table S1). These isolates were selected due to nutrient competition being their main mechanism for pathogen suppression, eliminating confounding effects from secondary metabolites^23^. We conducted *in vitro* resource utilization experiments to fully characterize the metabolic capacity of each strain, subsequently refining models to address apparent gaps in purely genomics-based models. These models were then used to predict pathogen invasion, as well as the associated substrate and metabolite features (Fig. 1B). Finally, we validated these predictions experimentally, offering key insights into how microbial substrate and metabolite features contribute to pathogen suppression.

**Fig. 1.**
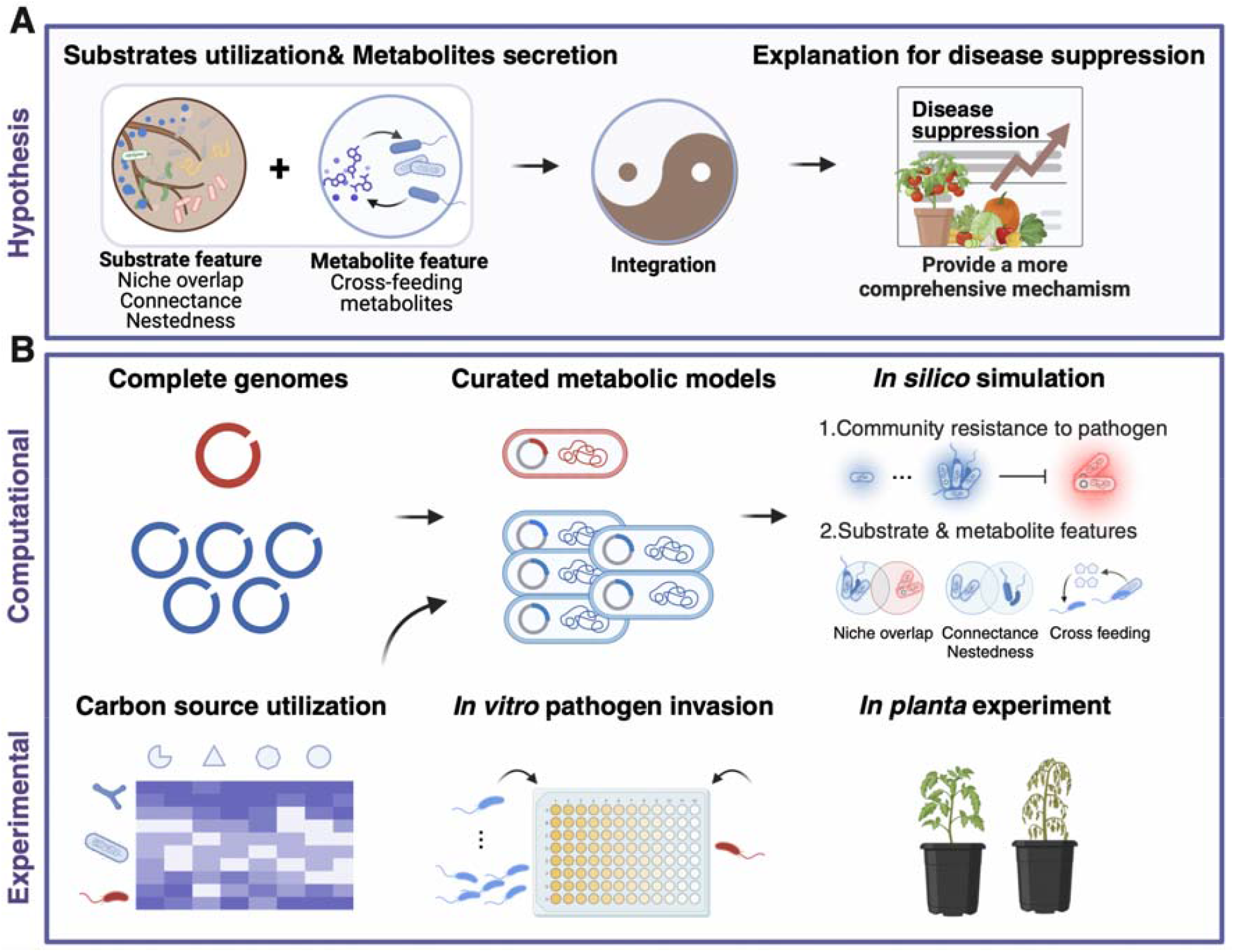
Hypothesis and the workflow. (**A**) Hypothesis scenarios. Plant roots release abundant carbon substrates that attract both native species and pathogens. The resident community can suppress invading pathogens by outcompeting them for nutrient substrate. However, the role of microbial metabolites in pathogen suppression is not well understood. We hypothesize that considering both substrate and metabolite features offers a more comprehensive understanding of disease suppression. In particular, microbial cross-feeding, although less explored, is likely to play a significant role in enhancing community resistance to pathogen invasion. (**B**) Experimentally and computationally guided workflow. Genome-scale metabolic models were constructed for both the pathogen and the resident community, based on genomic data and carbon source utilization profiles. Flux balance analysis was employed to simulate pathogen invasion and metabolic features across different community configurations and carbon source environments. These *in silico* predictions were then validated through *in vitro* and *in planta* experiments, offering key insights into the mechanisms underlying microbial community-mediated pathogen suppression.

## Results

### *In silico* simulation of pathogen invasion and metabolic features

We initially conducted *in vitro* carbon source utilization assays using 48 environmentally relevant substrates to characterize the metabolic capacity of each strain (Fig. 2A and Table S2). Draft models based solely on genomic data showed limited accuracy in predicting substrate utilization profiles, due to incomplete genome annotations and missing key metabolic pathways, which constrained their predictive power^20^. To address this, we curated genome-scale models by incorporating experimental carbon source utilization data to fill these gaps for each growth-supporting carbon source. This refinement significantly improved the accuracy of substrate utilization predictions, with a mean improvement of 52% (Student’s paired *t* test: *t* = -5.1, *df* = 5.0, *P* = 0.004, Fig. 2B). Next, we employed these models to simulate community resistance to pathogen invasion across a wide range of community and nutritional environments. This involved 620 pathogen invasion tests spanning 31 SynCom combinations (covering all possible species richness and community compositions, Table S3) and 20 randomly selected carbon source combinations (Table S4). Notably, our results demonstrated strong predictive capabilities of SynComs in pathogen suppression, achieving an F1 score of 96% (Fig. 2C). We further used genome-scale models to predict substrate and metabolite features. To represent substrate features, we included three variables— nestedness, connectance, and niche overlap—based on our previous work^7^. Niche overlap reflects substrate competition between native species and invader, nestedness and connectance describes substrate competition within native species. Our findings revealed a strong correlation between predicted and observed substrate features (*R*^*2*^ > 0.85, Fig. 2D), enabling us to incorporate the observed substrate features into subsequent analyses. For metabolite features, we used community models within a consistent nutritional environment to predict three variables—total produced metabolites, extracellular metabolites, and cross-feeding metabolites—to capture the metabolite exchanges during pathogen invasion tests (Fig. 2E). The number of these metabolites effectively limit pathogen invasion both *in vitro* and *in planta* (Fig. 3A-B), suggesting the importance of microbial metabolite in pathogen suppression. During microbial cross-feeding, strain QL-A2 displayed the highest frequency of absorbing metabolites (*ANOVA, F*_*4,95*_ = 67.1, *P* < 0.001), while strain QL-140 showed the highest frequency (*ANOVA, F*_*4,95*_ = 284.9, *P* < 0.001) of producing metabolites (Fig. S1).

**Fig. 2.**
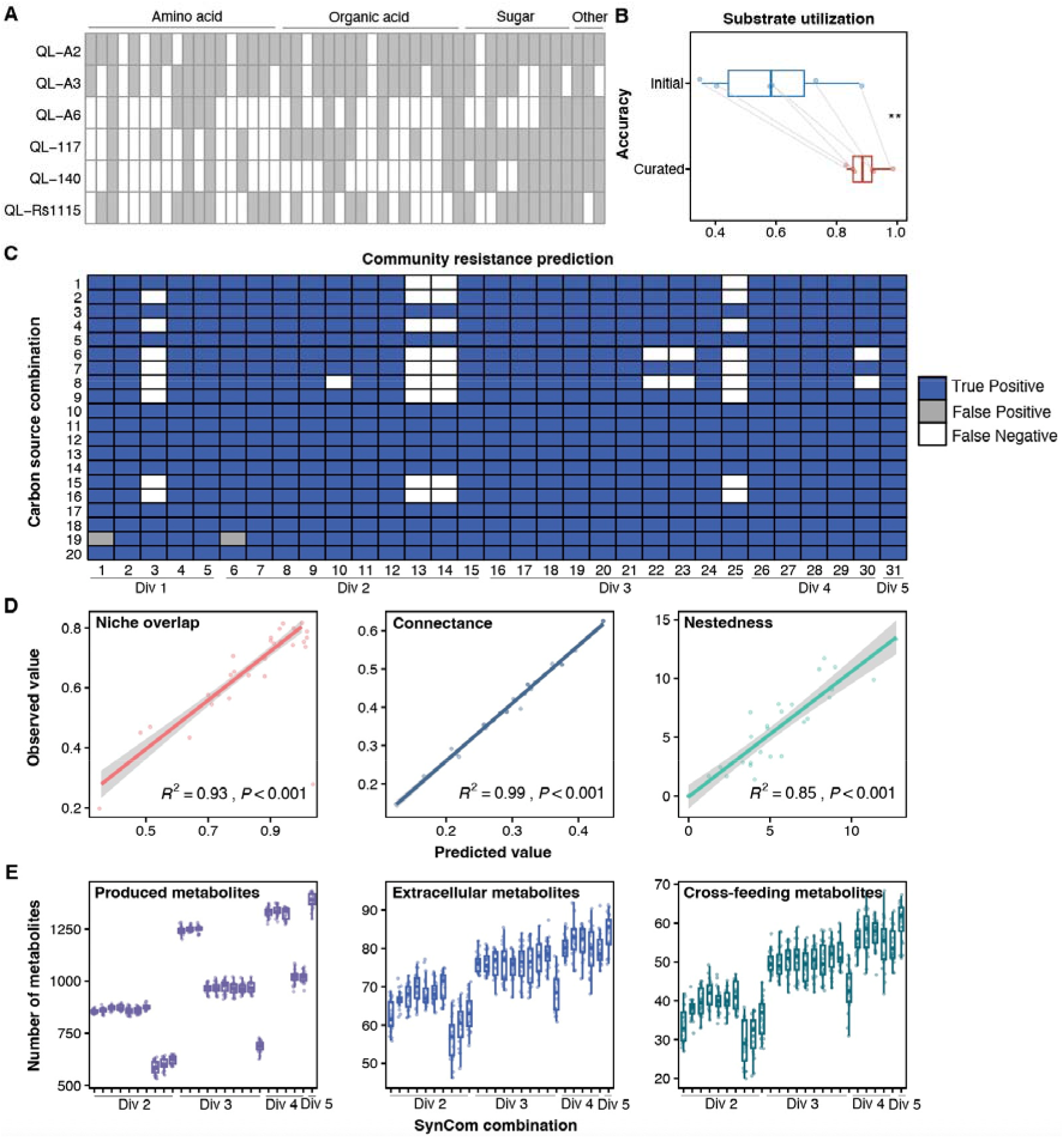
Modeling predictions for pathogen invasion and metabolic features. (**A**) Experimental carbon source utilization profiles of 48 common resources in the rhizosphere for each strain. The fill color of the boxes indicates whether growth occurred (grey) or did not occur (white) for each carbon resource. (**B**) Comparison of prediction accuracy for carbon source utilization between solely genomic-based models and curated models. Curated models were constructed through gap-filling based on the experimental carbon source utilization profiles of each strain. *P* values were calculated using pairwise Student’s *t*-tests (\**P* < 0.05, \*\**P* < 0.01). (**C**) Prediction accuracy matrix of putative interactions between 31 SynCom communities and the pathogen under 20 randomly selected carbon source combinations. Interactions are inferred by comparing pathogen growth in coculture versus monoculture. (**D**) Linear regression between predicted substrate features and observed substrate features. The *R*^*2*^ and *P* values denote the goodness of fit and significance level of linear regression, respectively. (**E**) Prediction of the number of total produced metabolites, extracellular metabolites, and cross-feeding metabolites under the same community and nutritional conditions as the *in vitro* pathogen invasion experiments. Only SynComs with a richness greater than 1 were included, as cross-feeding requires at least two microbial members for effective interaction. The dots in the boxplot represent the number of each metabolite under 20 carbon resource combinations.

**Fig. 3.**
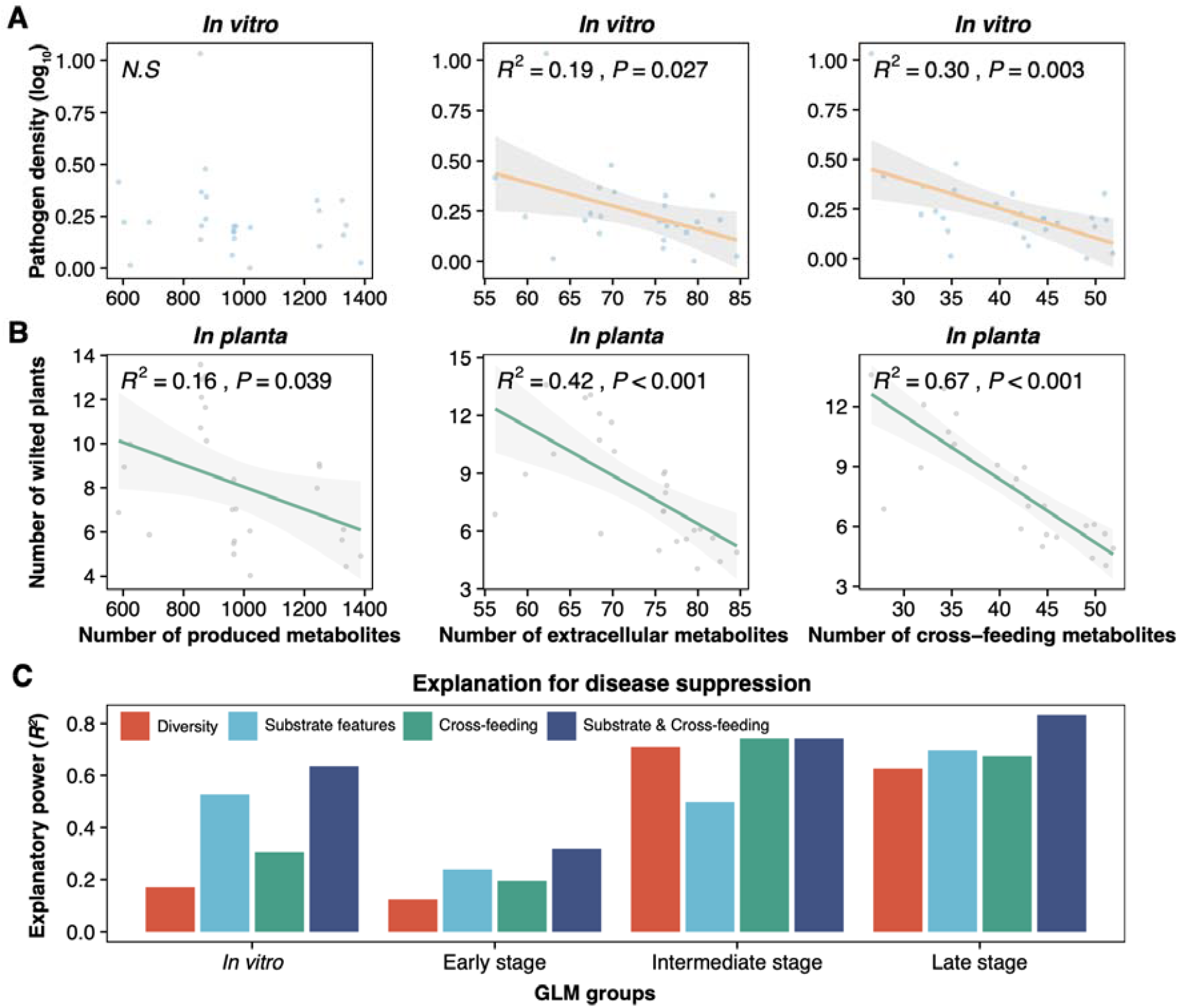
Impact of considering both substrate and metabolite features on explaining pathogen suppression. (**A**) Linear regression between the number of predicted metabolites (total produced, extracellular, and cross-feeding) and measured pathogen density. Only SynCom with a richness greater than 1 were included, as effective cross-feeding requires the presence of at least two microbial members for interaction. The dots represent the average number of metabolites in 20 different carbon resource combinations. (**B**) Linear regression between predicted metabolites (total produced, extracellular, and cross-feeding) and disease severity. Disease severity is measured by the number of wilted plants. (**C**) Comparison of the explanatory power for pathogen suppression between models considering both substrate and metabolite features versus those considering only community diversity, substrate features, and cross-feeding, based on generalized linear models (GLMs) results. Explanatory power is quantified by *R*^*2*^ values obtained from GLMs analyses.

### Considering both substrate and metabolite features improves explanation of pathogen suppression

In addition to *in vitro* pathogen invasion experiments, we extended our research to a greenhouse setting using tomato plants grown in sterile planting substrate, applying the same 31 SynCom treatments used in the *in vitro* assays. By combing data from both the *in vitro* and *in planta* experiments, we aimed to determine whether incorporating substrate and metabolite features could more effectively explain community resistance to pathogen invasion. Given the crucial role of cross-feeding metabolites in explaining community resistance to pathogen invasion, as observed across all metabolite features in both *in vitro* and *in planta* experiments (*in vitro*: *R*^*2*^=0.30, *in planta*: *R*^*2*^= 0.67, Fig. 3A-B), we incorporated cross-feeding metabolites with substrate features—nestedness, connectance, and niche overlap and employed generalized linear models (GLMs) to explore the explanation for pathogen suppression (Table 1). Our results revealed that considering both substrate and metabolite features offered a more comprehensive explanation for pathogen suppression than using community diversity, substrate features, or cross-feeding individually (Table 1 and Fig. 3C). This integration resulted in an average increase in explanatory power of 54.96%, 28.99%, and 31.98%, respectively (Fig. 3C). Remarkably, cross-feeding metabolites consistently emerged as significant predictors across all *in vitro* and *in planta* experiments when analyzed through stepwise regression (Table 1), highlighting the crucial role of cross-feeding in enhancing community resistance to pathogen invasion.

**Table 1.**
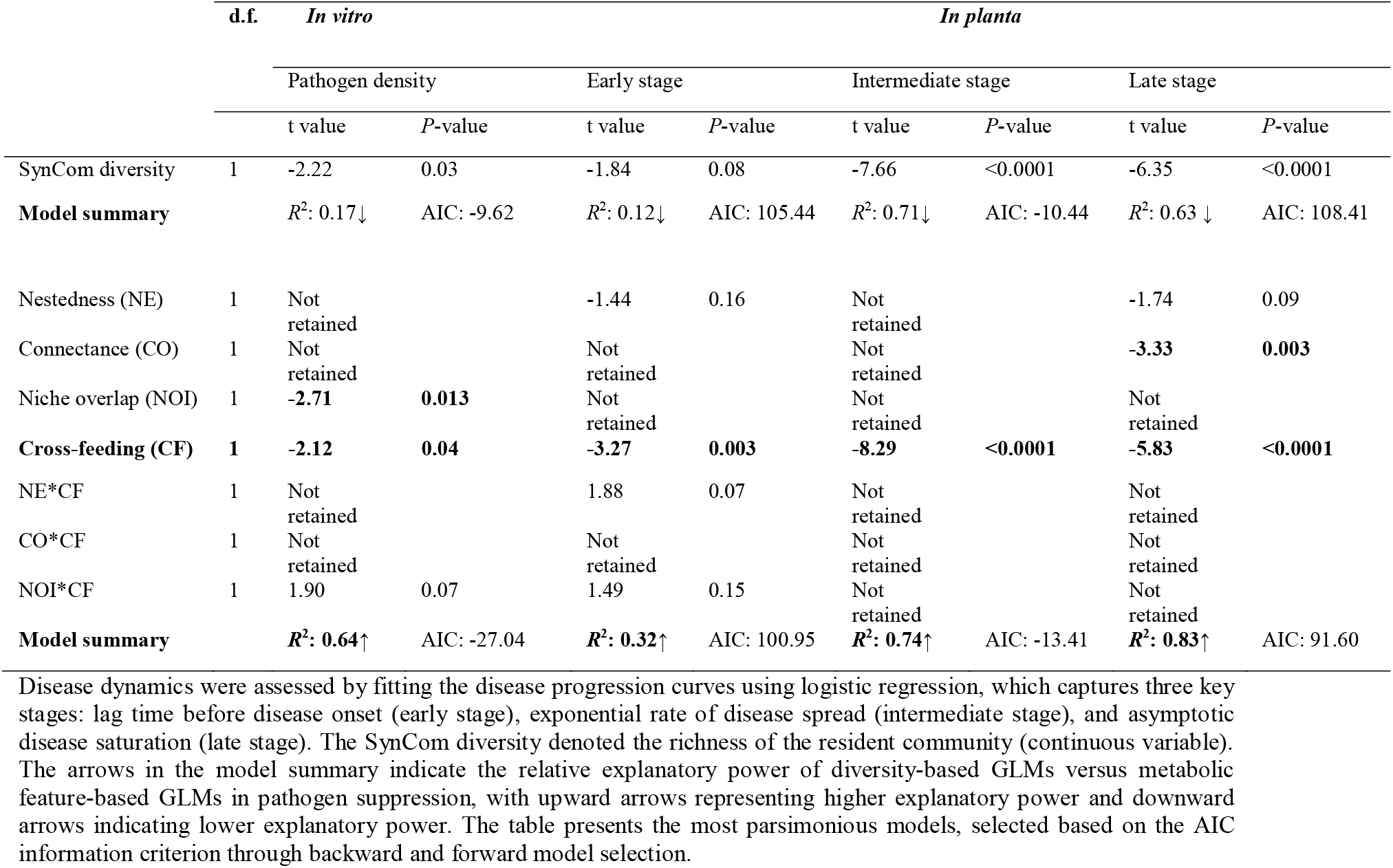
Comparison of the explanation of pathogen suppression between models that incorporate both substrate and metabolite features and those that consider diversity alone, based on generalized linear models (GLMs) results.

### Mechanism of cross-feeding metabolites in pathogen suppression

Given the important role of predicted cross-feeding metabolites in conferring community resistance to pathogen invasion observed in both *in vitro* and *in planta* experiments, we aimed to validate their function and elucidate the underlying mechanism. We first selected the SynCom with the highest species richness (richness = 5) and conducted *in silico* simulations by randomly selecting ten selected cross-feeding and ten non-crossfeeding metabolites from 48 carbon sources pool, repeating 99 times. Our simulations demonstrated that cross-feeding metabolites significantly suppressed pathogen density compared to non-crossfeeding metabolites (Student’s *t* test: *t* = -18.9, *df* = 119.8, *P* < 0.0001, Fig. 4A). This finding was further confirmed through *in vitro* experiments under same community and metabolite conditions (Student’s *t* test: *t* = -2.7, *df* = 193.5, *P* = 0.007, Fig. 4A). We hypothesize that pathogens have limited ability to utilize cross-feeding metabolites, while the native microbial community effectively leverages these metabolites to promote its growth (Fig. 4B). To test this hypothesis, we conducted an additional *in vitro* experiment comparing SynCom biomass and pathogen density under cross-feeding and non-crossfeeding metabolite conditions. The results showed significantly lower pathogen densities (Student’s *t* test: *t* = -28.9, *df* = 184.9, *P* < 0.0001, Fig. 4C) and higher SynCom growth (Student’s *t* test: *t* = 2.66, *df* = 184.1, *P* = 0.009, Fig. 4C) in the presence of cross-feeding metabolites. Furthermore, we observed a negative correlation between SynCom biomass and pathogen density in the presence of cross-feeding metabolites (Fig. S2). These findings confirm that pathogens are less effective at utilizing cross-feeding metabolites, while these metabolites enhance SynCom growth.

**Fig. 4.**
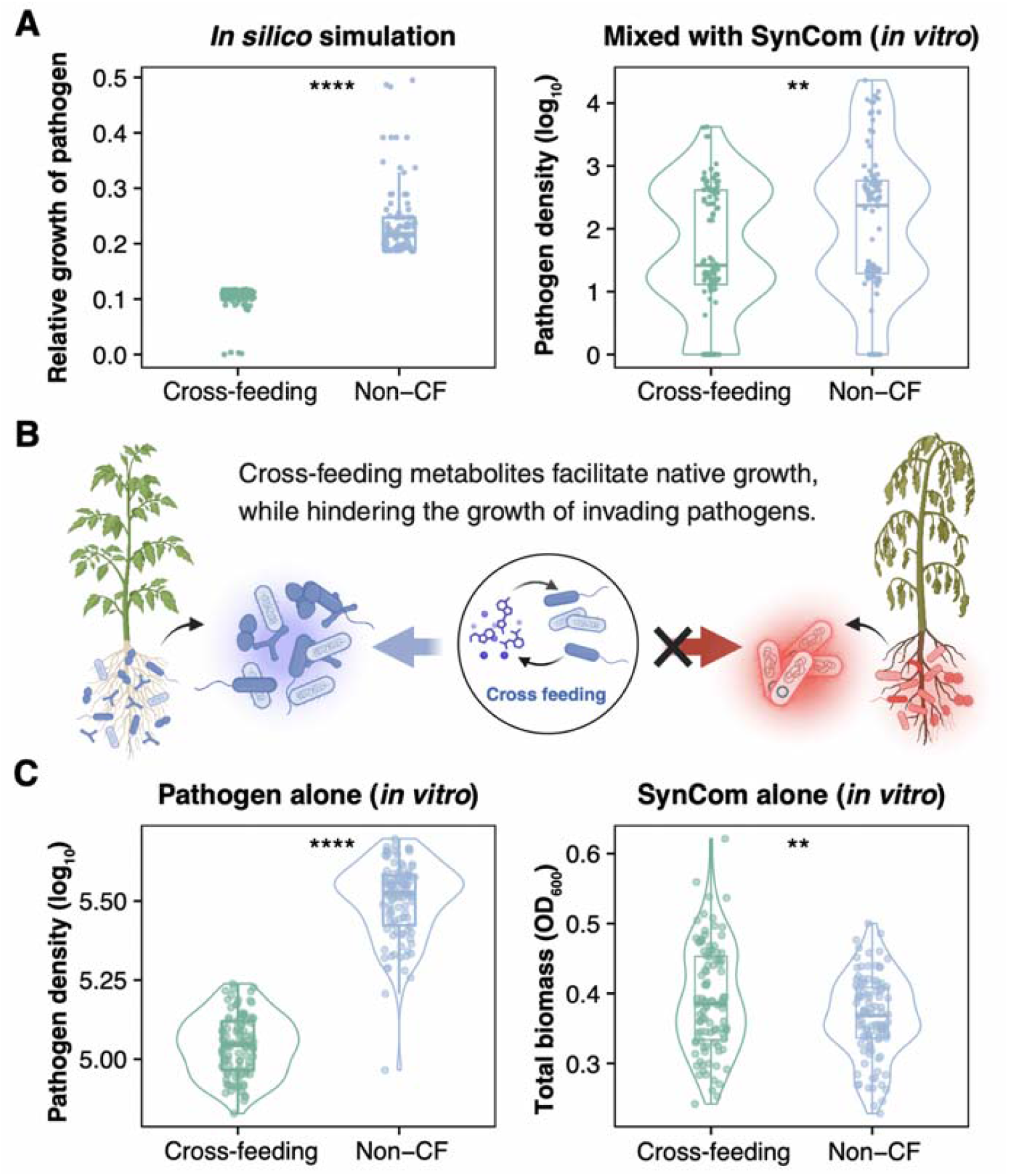
Experimental validation of the role cross-feeding metabolites in pathogen suppression. (**A**) Comparison of the pathogen growth and density with the addition of cross-feeding metabolites versus non-crossfeeding metabolites, as observed in both *in silico* simulation and *in vitro* validation. Non-CF denotes non-crossfeeding metabolites. The relative pathogen growth is defined as the ratio of pathogen growth to the resident community growth. (**B**) Schematic illustrating the hypothesis that pathogens are less efficient at utilizing cross-feeding metabolites, while these metabolites enhance the growth of native species. (**C**) Comparison of *in vitro* pathogen density and SynCom growth with the addition of cross-feeding metabolites versus non-crossfeeding metabolites. *P* values were calculated using Student’s *t*-tests (\**P* < 0.05; \*\**P* < 0.01; \*\*\**P* < 0.001; \*\*\*\**P* < 0.0001).

## Discussion

In this study, we investigated the utility of genome-scale metabolic models in predicting pathogen invasion across diverse microbial communities and nutritional environments, shedding light on the mechanisms underlying microbiome resistance against pathogen invasion. Our findings demonstrate the strong performance of genome-scale models in predicting community resistance to pathogen invasion, achieving a 96% F1 score across 620 invasion tests. This highlights the promising role of genome-scale models as a valuable tool for the targeted design of disease-resistant microbiomes, offering potential strategies to mitigate pathogen-related risks in microbial ecosystems. Moreover, our results indicated that considering both substrate and metabolite features offered a more robust explanation for microbiome resistance to pathogen invasion than community diversity, substrate feature, or cross-feeding alone. Notably, cross-feeding metabolites emerged as an a critical but often overlooked factor in conferring resistance against pathogen invasion. Subsequent *in vitro* experiments revealed the mechanism behind these metabolites, showing that pathogens struggle to utilize those cross-feeding metabolites, while these metabolites enhance the vigor and competitiveness of the resident community. Overall, our study not only emphasizes the crucial role of genome-scale models in predicting community resistance to pathogen invasion but also highlights the importance of considering substrate and metabolite features to better explaining pathogen suppression.

Differences in substrate utilization profiles between models based solely on genomic data and those informed by experimental data have been widely reported. This discrepancy is often attributed to inherent gaps and inaccuracies in genome annotations^24^, failing to fully capture the true metabolic capacities of strains^25,26^. When key metabolic pathways are absent or improperly annotated, strains struggle to utilize certain growth-supporting resources effectively, leading to incorrect predictions about their biomass production under different environmental conditions^27^. Therefore, *in vitro* carbon source utilization profiles were used for gap-filling to refine the models, which allowed us to create more robust and reliable models capable of simulating and predicting the microbial interactions in various nutritional environments^26^. In addition to gap-filling based on the carbon source screen, the strains in our study exhibit metabolic competition without direct antagonism may provide a solid foundation for exploring their interactions and the underlying metabolic mechanisms based on genome-scale models^28^. Moreover, future research could integrate more experimental and multi-omics data, such as growth curves, transcriptomics, and metabolomics, which allow for a more detailed understanding of gene expression and metabolite production, thereby improving the quantitative predictive power^29^. Future research could include a larger and more diverse community to explore the predictive accuracy in pathogen invasions and the design of beneficial communities targeted to suppress specific pathogens more effectively.

Cross-feeding plays a pivotal role in pathogen suppression by promoting cooperation and maintaining functional diversity within microbial communities^30,31^. It facilitates the exchange and recycling of essential metabolites among microbes and ensures a relatively balanced ecosystem^32^. This collaborative resource sharing through cross-feeding is particularly important for creating a robust defense against invading pathogens, as it enhances the community’s collective metabolic capabilities^33^. In healthy microbiomes, cross-feeding networks contribute to metabolic redundancy and cooperation, where different microbes contribute to the overall fitness and stability of the community^34^. However, when cross-feeding networks break down, as seen in dysbiotic states, the balance of resource sharing is disrupted^35^. Dysbiosis often results in a loss of functional diversity, where certain microbes are outcompeted or eliminated, leaving gaps in resource utilization. Pathogens can exploit these gaps, utilizing unclaimed nutrients and proliferating in the absence of competition^11^.

From an evolutionary perspective, cross-feeding reflects processes such as Black Queen evolution^36^, where some species lose the ability to produce certain metabolites and instead rely on other members of the community to provide them. In this scenario, cross-feeding is not just a cooperative behavior but an evolutionary necessity, where metabolic functions are distributed among different species^37^. As microbial species evolve to specialize and share metabolic functions, they form a tightly interconnected community where each member plays a critical role in maintaining overall function^38^. The co-adaptive nature of cross-feeding helps explain why disruptions in these networks, such as those caused by environmental changes or antibiotic use, can lead to pathogen susceptibility. In the absence of effective cross-feeding, pathogens can more easily invade and disrupt the functional balance of microbiomes, but strong cross-feeding networks acts as a metabolic buffer, enhancing the community resistance to pathogens^34^.

In summary, our study highlights the utility of genome-scale models in predicting community resistance to pathogen invasion across diverse nutritional environments and in elucidating the underlying metabolic mechanisms determining this resistance. This paves the way for the targeted design of disease-resistant microbiomes. By considering both substrate and metabolite features, we offer a more comprehensive understanding of the factors contributing to microbiome resistance within diverse microbiomes. This work not only advances our understanding of microbial interactions but also offers practical applications for improving ecosystem health.

## Methods

### Bacterial strains

The phytopathogen used in this study was *Ralstonia solanacearum* QL-Rs1115, which was tagged with the pYC12-mCherry plasmid^39^. Additionally, we utilized five nonvirulent *Ralstonia* strains (*Ralstonia mannitolilytica* QL-A2, *Ralstonia mannitolilytica* QL-A3, *Ralstonia pickettii* QL-A6, *Ralstonia* sp. QL-117, *Ralstonia pickettii* QL-140) as commensal species^7^. Notably, no antagonistic interactions were detected either among the commensal species or between the commensal species and the phytopathogen^23^. All strains were routinely maintained at -80 □with glycerol (final concentration of 20%). Further details on the bacterial strains and plasmid were listed in Table S1.

### Carbon source utilization experiment

Each strain was cultured in Nutrient Broth (NB) medium (composed of glucose 10.0 g L^-1^, peptone 5.0 g L^-1^, yeast extract 0.5g L^-1^, beef extract 3.0g L^-1^, pH 7.0) at 30 □until reaching the exponential growth phase. Subsequently, cells were harvested by centrifugation at 6000 rpm for 6 minutes, washed three times with 0.85% NaCl buffer, and resuspended in OS minimal medium^40^ to an optical density at 600 nm (OD_600_) of 0.1. The bacterial suspension was then transferred to 150 μL OS minimal medium supplemented with a specific resource in individual wells of 96-well plates, with six replicates per resource condition. We used 48 commonly detected resources from tomato rhizosphere to represent environmental substrates (Table S2). OS minimal medium without any resource was used as the negative control. After 48 h growth at 30 □with agitation, optical density of the single strain was measured using a SpectraMax M5 Plate reader (Molecular Devices, Sunnyvale, CA, USA). Wells with an OD_600_ > 0.05 were considered as positive growth on the respective resource.

### *In vitro* pathogen invasion experiment

Prior to the experiment, a single colony of each strain was selected and grown overnight on Nutrient Agar (NA, NB medium with agar 18 g L^-1^). The bacterial cultures were subsequently washed three times with 0.85% NaCl buffer and adjusted to an OD_600_ of ∼0.5, approximately 10^7^ CFU ml^-1^. We systematically assembled 31 microbial communities covering all possible species diversity and community compositions (Table S3), following substitutive designs^41^. Then, each SynCom was established with an initial total bacterial density of 10^7^ CFU mL^-1^ to ensure consistency across all combinations, and the ratios of strains in SynComs with richness above 2 were equal. Next, SynComs were inoculated into 20 different resource combinations to simulate various nutritional environments (Table S4), resulting in a total of 620 pathogen invasion experiments (31 communities x 20 nutritional environments, Table S5). Specifically, each SynCom consisted of OS minimal medium supplemented with a random mixture of 20 resources selected from a total pool of 48 resources. The SynComs were then exposed to invasion by the mCherry-tagged *R. solanacearum* QL-Rs1115 strain (at a concentration of 10^5^ cells per ml) in 96-well plates (200 μl per well) at 30 □with agitation, while SynComs grown alone in the absence of the pathogen served as negative controls. SynCom growth was assessed after 48 hours based on OD_600_, while pathogen growth was evaluated based on mCherry fluorescence signal (excitation: 587 nm, emission: 610 nm). Next, we adjusted the red fluorescent protein signals (RFP) by subtracting the RFP measured in the negative control (absence of the pathogen) from the RFP measured in the presence of the pathogen. We then calculated pathogen density using the formula log_10_ (RFP/OD_600_). A successful invasion was defined as a scenario where the pathogen density was greater than 0 and the RFP of the treatment with the pathogen exceeded the average RFP value of the negative control without the pathogen. Only communities that experienced successful invasion were considered for further analysis^7^. An interaction was defined as suppression if pathogen density in coculture was less than 90% of that in monoculture. Conversely, it was defined as facilitation if these values were more than 110% of those in monoculture^20^.

### Greenhouse experiment

A 42-day greenhouse experiment was conducted to investigate the inhibitory effect of 31 SynComs against the pathogen. Consistent with our *in vitro* experiments, we utilized the same 31 SymComs with two replicates each, with positive controls containing only the pathogen (three replicates), and negative controls without any bacteria (three replicates). Tomato seeds (Lycopersicon esculentum, cultivar ‘Jiangshu’) were surface-sterilized and germinated on a water agar plate for 3 days before being transferred to seedling plates containing cobalt-60-sterilized seedling substrate sourced from the Huainong Institute of Soil and Fertilizer, Huaian, China. At the three-leaf stage (11 days after seeding), tomato plants were then transplanted into seedling trays containing the same seedling substrate and inoculated with SynCom via drenching, resulting in an ending concentration of 10^8^ CFU/g soil^42^. The invasion experiment started after 1 week by inoculating the pathogen, resulting in a final concentration of 10^6^ CFU/g soil. Throughout the experiment, the greenhouse maintained a temperature range of 25-35 °C and was regularly watered with sterile water to ensure optimal tomato growth. To mitigate bias, the seedling pots were randomly rearranged every 2 days. Daily monitoring of the number of wilted seedlings per plate from day 17 after transplantation. Given the inability to directly quantify available rhizosphere carbon resources, we assumed that all 48 resources utilized in our *in vitro* experiments were available in the rhizosphere. Disease dynamics was evaluated using a logistic growth curve that tracked the percentage of wilted plants as a function of time throughout the experiment’s duration. This curve fitting yielded three variables: the lag phase (days after pathogen inoculation) before disease onset, the intrinsic rate of increase of wilted plants (day^-1^), and the asymptotic disease prevalence (% of wilted plants), which are respectively referred to as the early, intermediate, and late stages of infection.

### DNA extraction and genome sequencing

The genomic DNA extraction of six strains was performed using a Bacteria DNA Kit (OMEGA) following the manufacturer’s instructions, with subsequent quality control of the purified DNA samples. Genomic DNA was quantified using a TBS-380 fluorometer (Turner BioSystems Inc., Sunnyvale, CA). High-quality DNA samples (OD_260/280_=1.8∼2.0, >6 μg) were used to construct fragment libraries. The genomes of the strains were sequenced employing a combination of PacBio RS and Illumina sequencing platforms. For Pacific sequencing, 20 kb insert whole-genome shotgun libraries were prepared and sequenced on a Pacific Biosciences RS instrument using standard protocols. An aliquot of 8 μg DNA was spun in a Covaris g-TUBE (Covaris, MA) at 6,000 RPM for 60 seconds using an Eppendorf 5424 centrifuge (Eppendorf, NY). DNA fragments were then purified, end-repaired, and ligated with SMRTbell sequencing adapters following the manufacturer’s recommendations (Pacific Biosciences, CA). Resulting sequencing libraries were purified three times using 0.45x volumes of Agencourt AMPure XP beads (Beckman Coulter Genomics, MA) following the manufacturer’s recommendations. For Illumina sequencing, at least 3 μg of genomic DNA was used for library construction. Paired-end libraries with insert sizes of approximately 400 bp were prepared following Illumina’s standard genomic DNA library preparation protocol. Purified genomic DNA was sheared into smaller fragments with a desired size by Covaris, and blunt ends were generated using T4 DNA polymerase. After adding an ‘A’ base to the 3’ end of the blunt phosphorylated DNA fragments, adapters were ligated to the ends of the DNA fragments. The desired fragments were purified through gel electrophoresis, then selectively enriched and amplified by PCR. The index tag could be introduced into the adapter at the PCR stage as appropriate, and a library quality test was performed. The qualified Illumina paired-end library was then used for Illumina NovaSeq 6000 sequencing (150 bp * 2, Shanghai BIOZERON Co., Ltd). The raw paired-end reads were trimmed and quality controlled by Trimmomatic^43^ with the parameters (SLIDINGWINDOW: 4:15 MINLEN:75). Clean data obtained from the quality control processes were used for further analysis. Illumina sequencing data was used to evaluate the complexity of the genome and correct the PacBio long reads. ABySS^44^ was used for genome assembly with multiple-Kmer parameters to obtain the optimal assembly results, and Canu^45^ was used to assemble the PacBio-corrected long reads. GapCloser^46^ was subsequently applied to fill up the remaining local inner gaps and correct the single base polymorphisms for the final assembly results. The quality assessment of these genomes was conducted using CheckM^47^, revealing an average completeness of 99.88% (Table S1).

### Construction of genome-scale metabolic models

The gapseq^48^ tool was employed to construct genome-scale metabolic models and perform gap-filling. This tool utilizes various data sources and a novel gap-filling procedure, enhancing the versatility of metabolic models for predicting physiological responses under various chemical growth conditions^48^. Initially, metabolic models were generated based on the assembled genomes of each strain. Subsequently, we compared the predicted carbon source utilization of each draft model with experimental data from carbon source utilization assays, and those carbon sources that the strain could utilize but were not accurately predicted by the model were identified. We then used gapseq to perform gap-filling on each graft model to generate a new set of biosynthesis and transport reactions, allowing the model to produce biomass from each growth-supporting carbon source identified in our experiments.

### Computing predicted pathogen growth in monoculture and coculture based on genome-scale models

We utilized COBRApy^49^ with the CPLEX solver v12.10 (IBM) in Python 3.7 to compute the pathogen growth in both monoculture and coculture conditions. Similar to our *in vitro* experiments, the simulated medium consisted of OS minimal medium and 20 resource combinations. Each resource combination consisted of 20 randomly picked resources out of 48 resources. The models were provided with nonlimiting amounts (*v*_max_ = 1000 mmol gDW^−1^ hour^−1^) of the simulated OS minimal medium components (Table S6), including essential carbon dioxide, metal ions, water, and nitrogen, sulfur, and phosphorus sources. Additionally, limiting amounts (*v*_max_ = 0.1 mmol gDW^−1^ hour^−1^) of the resources were supplied in the medium. For the SynCom models, individual strain models were integrated into a shared extracellular compartment^50,51^, with all metabolites, reactions, and genes specifically labeled, except those in the extracellular compartment. Biomass was set as the objective function for model growth simulations. To simulate an equal abundance of resources between monoculture and coculture conditions, the sum of biomass reactions was set as the objective to be optimized, and exchange reaction values were adjusted to match those of the models in monoculture. Additionally, the model was constrained to prevent the uptake of resources outside of those provided in the defined medium. For each pair or community, we simulated the biomass fluxes of the pathogen when grown alone and cocultured with the SynCom. Community resistance to pathogen invasion was assessed by comparing predicted pathogen growth in coculture to pathogen growth in monoculture. Resistance was defined as pathogen growth in coculture being less than 90% of that in monoculture, a threshold commonly employed to identify microbial interactions^20^.

### Computing metabolite features within the SynCom based on genome-scale models

The predicted numbers of produced metabolites, extracellular metabolites, and cross-feeding metabolites within the SynComs were calculated based on COBRApy^49^ with the CPLEX solver v12.10 (IBM) in Python 3.7. Metabolites produced by one strain and absorbed by another within the communities were identified as cross-feeding metabolites, which include both mutualistic and exploitative interactions^10^. Considering the cross-feeding requires the presence of two or more strains, we only included the SynComs with a richness at least 2 in our predictions of produced metabolites and extracellular metabolites. We used the same SynCom and nutrient environment in pathogen invasion tests to simulate metabolite exchanges within microbial community. For the SynCom models, individual strain models were integrated into a common extracellular compartment^50,51^, with all metabolites, reactions, and genes specifically labeled, except those in the extracellular compartment. Metabolites with labels containing “_e” were identified as extracellular metabolites.

### Additional *in vitro* validation experiment

To further investigate the role of cross-feeding metabolites in suppressing pathogen invasion, we conducted additional validation experiments based on *in silico* simulation and *in vitro* experiment. We began by selecting the resident community with the highest richness (richness = 5) to maximize their cross-feeding potential. From our pool of carbon sources, we randomly selected 10 cross-feeding metabolites and 10 non-crossfeeding metabolites to predict the pathogen growth when the resident community was cocultured with the pathogen *in silico*, repeating 99 times. Following the *in silico* conditions, we conducted an *in vitro* experiment using the same community and nutritional environments. We conducted three different treatments: (1) pathogen grown alone, (2) pathogen cocultured with the resident community, and (3) resident community cultured alone. We then measured the RFP and OD_600_ to evaluate pathogen density and SynCom biomass.

### Data analysis

Analysis of variance (ANOVA) were performed using the aov() function of the stats package. Linear regressions were used to explore the relationship between metabolic features and pathogen density. General linear models (GLMs) were employed to assess the explanatory power of metabolic features and community diversity in pathogen suppression. Due to potential correlations among metabolic features, we used stepwise model selection (step() function of the stats package) to identify the optimal GLMs. Student’s *t*-tests were used to test statistical significance between pairs of samples, where *P* value below 0.05 were considered statistically significant. If not stated otherwise, all data processing, statistical analyses, and graphical representation were performed in R 4.1.3 (www.r-project.org).

## Data availability

The genome sequences and metabolic models are available on Zenodo (10.5281/zenodo.14007949).

## Code availability

The scripts used for analysis are available on Zenodo (10.5281/zenodo.14008038).

## Acknowledgements

We thank Jiaheng Hou from Peking University for insightful discussions and suggestions. This research was funded by the National Natural Science Foundation of China (42325704, 42477125, and 42090060), the Fundamental Research Funds for the Central Universities (KYT2024001), the Natural Science Foundation of Jiangsu Province (BK20230102), the Jiangsu Carbon Peak & Carbon Neutrality Science and Technology Innovation Special Fund (BE2022423), and Postgraduate Research & Practice Innovation Program of Jiangsu Province (KYCX24_0970).

## Author contributions

Conceptualization: X.Y., T.Y. and Z.W.; Resources: T.Y., A.J., and Z.W.; Methodology: X.Y. T.Y., and A.J.; Data curation: X.Y., Z.Z., Y.Z., X.M., Y.G., N.W., and G.J.; Formal analysis: X.Y., Z.Y., and N.W.; Funding acquisition: T.Y., G.J, Y.C., Q.S., M.H.M. and Z.W.; Investigation: X.Y., Z.Z., Y.Z., X.M., Y.G., N.W., and G.J.; Project Administration: T.Y., Z.W., Y.C., and Q.S.; Supervision: T.Y., Z.W., Y.C., and Q.S.; Software: X.Y., T.Y.; Visualization: X.Y., T.Y., Y.Z., and A.J.; Writing—original draft: X.Y., T.Y. and A.J.; Writing—review & editing: T.Y., M.H.M., Z.W. and A.J.

## Declaration of interests

The authors declare no competing interests.

## Notes

### Competing Interest Statement

The authors have declared no competing interest.

### Summary of Updates

This version of the manuscript has been revised to update the following : Funding updated.

