## Supplementary files for "Substrate utilization and cross-feeding synergistically determine microbiome resistance to pathogen invasion" (from another version of this preprint: version 2, 2024-11-11)

### **S****upplementary Information**

*******Correspondence**

**This PDF file includes:**

**Table S1 to Table S6**

**Fig. S1 to Fig. S2**

| **Table S1 The basic information of strains and plasmid used in this study** | | |
| --- | --- | --- |
| Strain | Species | Genome completeness |
| QL-A2 | *Ralstonia mannitolilytica* | 100 |
| QL-A3 | *Ralstonia mannitolilytica* | 100 |
| QL-A6 | *Ralstonia pickettii* | 99.94 |
| QL-117 | *Ralstonia* sp. QL-117 | 99.94 |
| QL-140 | *Ralstonia pickettii* | 99.48 |
| QL-Rs1115 | *Ralstonia solanacearum* | 99.94 |
| Plasmid | PYC12-M | mCherry inserted in the pYC12, under control of the constitutive Ptac promotor, gmR |

**Table S2** **List of resources included in the metabolic screen and utilization profiles for each strain**

| Resource | Type | Name | QL-A2 | QL-A3 | QL-A6 | QL-117 | QL-140 | QL-Rs1115 |
| --- | --- | --- | --- | --- | --- | --- | --- | --- |
| L-Alanine | Amino acid | lala | 0 | 0 | 1 | 1 | 1 | 1 |
| beta-Alanine | Amino acid | bala | 0 | 1 | 1 | 1 | 1 | 0 |
| L-Arginine | Amino acid | arg | 0 | 0 | 0 | 0 | 0 | 0 |
| L-Asparagine | Amino acid | asn | 1 | 1 | 1 | 1 | 1 | 1 |
| Citrulline | Amino acid | cin | 0 | 0 | 1 | 1 | 1 | 1 |
| L-Glutamine | Amino acid | gln | 1 | 1 | 1 | 1 | 1 | 1 |
| L-Glycine | Amino acid | gly | 0 | 0 | 1 | 0 | 1 | 0 |
| L-Histidine | Amino acid | his | 0 | 1 | 1 | 1 | 1 | 1 |
| Isoleucine | Amino acid | iso | 1 | 0 | 0 | 1 | 1 | 0 |
| L-Lysine | Amino acid | lys | 0 | 0 | 0 | 0 | 0 | 0 |
| L-Leucine | Amino acid | leu | 0 | 0 | 0 | 1 | 1 | 0 |
| L-Methionine | Amino acid | met | 0 | 0 | 0 | 0 | 0 | 0 |
| L-Phenyalanine | Amino acid | phe | 0 | 0 | 1 | 1 | 1 | 1 |
| L-Proline | Amino acid | pro | 1 | 1 | 1 | 1 | 1 | 1 |
| L-Serine | Amino acid | ser | 0 | 0 | 1 | 0 | 0 | 1 |
| L-Threonine | Amino acid | thr | 0 | 0 | 1 | 1 | 1 | 0 |
| L-Tryptophan | Amino acid | try | 0 | 0 | 1 | 1 | 1 | 0 |
| L-Valine | Amino acid | val | 0 | 1 | 1 | 1 | 1 | 0 |
| Acetic acid | Organic acid | ace | 0 | 0 | 1 | 0 | 1 | 1 |
| Ascorbic acid | Organic acid | asc | 0 | 0 | 0 | 0 | 1 | 0 |
| gamma aminobutyric acid | Organic acid | ami | 1 | 1 | 1 | 0 | 1 | 1 |
| Citric acid | Organic acid | cit | 0 | 0 | 1 | 0 | 1 | 1 |
| Formic acid | Organic acid | for | 0 | 0 | 0 | 0 | 0 | 0 |
| Galacturonic acid | Organic acid | galac | 0 | 0 | 1 | 0 | 0 | 1 |
| Glutaric acid | Organic acid | glut | 0 | 0 | 0 | 0 | 1 | 0 |
| Glycolic acid | Organic acid | glyc | 0 | 0 | 1 | 1 | 1 | 0 |
| Lactic acid | Organic acid | lac | 1 | 1 | 1 | 1 | 1 | 1 |
| Maleic acid | Organic acid | male | 0 | 0 | 0 | 0 | 1 | 0 |
| Malic acid | Organic acid | mal | 0 | 0 | 1 | 1 | 1 | 1 |
| Malonic acid | Organic acid | malon | 0 | 0 | 1 | 1 | 1 | 0 |
| 2-Oxoglutaric | Organic acid | oxo | 0 | 0 | 1 | 0 | 1 | 1 |
| Pyruvic acid | Organic acid | pyr | 0 | 1 | 1 | 1 | 1 | 1 |
| Succinic acid | Organic acid | succ | 0 | 0 | 1 | 1 | 1 | 1 |
| Tartaric acid | Organic acid | tar | 0 | 0 | 0 | 0 | 0 | 1 |
| Oxalic acid | Organic acid | oxa | 0 | 0 | 0 | 0 | 0 | 0 |
| Fructose | Sugar | fruc | 0 | 0 | 1 | 0 | 1 | 0 |
| Glucose | Sugar | glu | 1 | 1 | 1 | 0 | 0 | 1 |
| Sucrose | Sugar | sucr | 0 | 0 | 0 | 0 | 0 | 1 |
| Maltose | Sugar | malt | 0 | 0 | 1 | 0 | 1 | 0 |
| L-Arabinose | Sugar | ara | 0 | 0 | 1 | 0 | 1 | 0 |
| D-Galactose | Sugar | gal | 0 | 1 | 1 | 0 | 0 | 0 |
| D-Mannose | Sugar | man | 0 | 1 | 1 | 0 | 0 | 0 |
| D-Xylose | Sugar | xyl | 0 | 0 | 0 | 0 | 0 | 0 |
| D-Ribose | Sugar | rib | 0 | 0 | 0 | 0 | 0 | 0 |
| D-Mannitol | Sugar | mann | 0 | 1 | 0 | 0 | 0 | 0 |
| Ethanolamine | Other | eth | 0 | 0 | 0 | 0 | 0 | 0 |
| L-Inositol | Other | mino | 0 | 0 | 0 | 0 | 0 | 1 |
| Inosine | Other | ino | 0 | 1 | 0 | 0 | 0 | 0 |

| **Table S3 The composition of 31 SynComs used in the study** | | | | | | | |
| --- | --- | --- | --- | --- | --- | --- | --- |
| No. | QL-A2 | QL-A3 | QL-A6 | QL-117 | QL-140 | QL-Rs1115 | Diversity |
| 1 | 1 | 0 | 0 | 0 | 0 | 1 | 1 |
| 2 | 0 | 1 | 0 | 0 | 0 | 1 | 1 |
| 3 | 0 | 0 | 1 | 0 | 0 | 1 | 1 |
| 4 | 0 | 0 | 0 | 1 | 0 | 1 | 1 |
| 5 | 0 | 0 | 0 | 0 | 1 | 1 | 1 |
| 6 | 1 | 1 | 0 | 0 | 0 | 1 | 2 |
| 7 | 1 | 0 | 1 | 0 | 0 | 1 | 2 |
| 8 | 1 | 0 | 0 | 1 | 0 | 1 | 2 |
| 9 | 1 | 0 | 0 | 0 | 1 | 1 | 2 |
| 10 | 0 | 1 | 1 | 0 | 0 | 1 | 2 |
| 11 | 0 | 1 | 0 | 1 | 0 | 1 | 2 |
| 12 | 0 | 1 | 0 | 0 | 1 | 1 | 2 |
| 13 | 0 | 0 | 1 | 1 | 0 | 1 | 2 |
| 14 | 0 | 0 | 1 | 0 | 1 | 1 | 2 |
| 15 | 0 | 0 | 0 | 1 | 1 | 1 | 2 |
| 16 | 1 | 1 | 1 | 0 | 0 | 1 | 3 |
| 17 | 1 | 1 | 0 | 1 | 0 | 1 | 3 |
| 18 | 1 | 1 | 0 | 0 | 1 | 1 | 3 |
| 19 | 1 | 0 | 1 | 1 | 0 | 1 | 3 |
| 20 | 1 | 0 | 1 | 0 | 1 | 1 | 3 |
| 21 | 1 | 0 | 0 | 1 | 1 | 1 | 3 |
| 22 | 0 | 1 | 1 | 1 | 0 | 1 | 3 |
| 23 | 0 | 1 | 1 | 0 | 1 | 1 | 3 |
| 24 | 0 | 1 | 0 | 1 | 1 | 1 | 3 |
| 25 | 0 | 0 | 1 | 1 | 1 | 1 | 3 |
| 26 | 1 | 1 | 1 | 1 | 0 | 1 | 4 |
| 27 | 1 | 1 | 1 | 0 | 1 | 1 | 4 |
| 28 | 1 | 1 | 0 | 1 | 1 | 1 | 4 |
| 29 | 1 | 0 | 1 | 1 | 1 | 1 | 4 |
| 30 | 0 | 1 | 1 | 1 | 1 | 1 | 4 |
| 31 | 1 | 1 | 1 | 1 | 1 | 1 | 5 |

“1” denotes to the presence of the species in the community and “0” denotes to the absence of species in the community.

| Com 1 | Com 2 | Com 3 | Com 4 | Com 5 | Com 6 | Com 7 | Com 8 | Com 9 | Com 10 | Com 11 | Com 12 | Com 13 | Com 14 | Com 15 | Com 16 | Com 17 | Com 18 | Com 19 | Com 20 |
| --- | --- | --- | --- | --- | --- | --- | --- | --- | --- | --- | --- | --- | --- | --- | --- | --- | --- | --- | --- |
| lala | bala | asc | cit | glu | glyc | male | malon | mino | pro | pyr | ser | tar | try | val | ara | man | xyl | rib | mann |
| bala | asc | asn | cin | for | galac | glu | glyc | leu | mal | met | oxo | pyr | tar | try | malt | gal | rib | mann | ino |
| lala | arg | asn | ami | cin | fruc | galac | iso | lac | lys | male | malon | phe | pro | try | val | man | xyl | rib | oxa |
| ace | lala | asn | ami | fruc | gln | glut | lys | leu | mal | malon | mino | oxo | succ | sucr | try | gal | man | rib | mann |
| ace | bala | arg | asc | cit | fruc | galac | gln | his | lac | male | malon | phe | succ | sucr | thr | man | xyl | rib | oxa |
| lala | asc | cit | gly | iso | lac | lys | male | mal | malon | mino | phe | pyr | ser | succ | sucr | tar | val | malt | mann |
| lala | bala | asc | asn | ami | for | glu | gly | lac | male | oxo | pro | pyr | sucr | ara | man | rib | mann | ino | oxa |
| ace | bala | asn | cit | cin | gln | glut | iso | male | mal | malon | met | phe | succ | sucr | tar | try | ara | xyl | oxa |
| lala | ami | cit | cin | for | his | mal | malon | met | oxo | pro | succ | thr | val | gal | man | xyl | mann | ino | oxa |
| ace | arg | asn | ami | cit | cin | galac | glu | gly | iso | leu | male | mino | phe | pro | pyr | succ | thr | malt | rib |
| ace | arg | asc | cit | for | gln | gly | iso | lys | leu | malon | mino | pro | pyr | ser | tar | try | gal | xyl | ino |
| ace | arg | asc | cin | for | galac | glu | gln | male | mino | phe | ser | succ | sucr | tar | gal | man | xyl | rib | ino |
| ace | arg | asn | ami | fruc | galac | gln | iso | lac | lys | leu | malon | oxo | phe | tar | thr | val | ara | rib | mann |
| ace | bala | arg | asn | cin | eth | for | gln | glyc | iso | male | phe | pyr | ser | tar | try | malt | ara | xyl | oxa |
| lala | bala | ami | for | fruc | gln | glut | gly | glyc | his | lac | mal | met | ser | sucr | tar | malt | ara | rib | ino |
| lala | bala | asc | cin | for | gln | gly | his | lys | malon | met | succ | sucr | tar | val | malt | man | xyl | rib | ino |
| ace | arg | asn | cin | eth | galac | glu | gly | his | lys | mal | mino | oxo | pro | pyr | try | ara | man | rib | oxa |
| lala | bala | arg | cit | fruc | glyc | iso | lys | male | mal | met | mino | oxo | tar | val | malt | rib | mann | ino | oxa |
| ace | lala | bala | arg | asc | cit | for | glu | iso | lys | leu | mal | mino | pyr | succ | try | malt | ara | gal | xyl |
| ace | arg | ami | cit | cin | galac | glu | gln | lys | leu | male | mal | met | oxo | pro | sucr | ara | xyl | rib | mann |

**Table S4 List of 20 resource combinations consisting of 20 resources**

| **Table S5 Information of *in vitro* pathogen invasion assay** | | | | | | | | |
| --- | --- | --- | --- | --- | --- | --- | --- | --- |
| Carbon.com | SynCom | A2 | A3 | A6 | QL117 | QL140 | Rs | div |
| 1 | 1 | 1 | 0 | 0 | 0 | 0 | 1 | 1 |
| 2 | 1 | 1 | 0 | 0 | 0 | 0 | 1 | 1 |
| 3 | 1 | 1 | 0 | 0 | 0 | 0 | 1 | 1 |
| 4 | 1 | 1 | 0 | 0 | 0 | 0 | 1 | 1 |
| 5 | 1 | 1 | 0 | 0 | 0 | 0 | 1 | 1 |
| 6 | 1 | 1 | 0 | 0 | 0 | 0 | 1 | 1 |
| 7 | 1 | 1 | 0 | 0 | 0 | 0 | 1 | 1 |
| 8 | 1 | 1 | 0 | 0 | 0 | 0 | 1 | 1 |
| 9 | 1 | 1 | 0 | 0 | 0 | 0 | 1 | 1 |
| 10 | 1 | 1 | 0 | 0 | 0 | 0 | 1 | 1 |
| 11 | 1 | 1 | 0 | 0 | 0 | 0 | 1 | 1 |
| 12 | 1 | 1 | 0 | 0 | 0 | 0 | 1 | 1 |
| 13 | 1 | 1 | 0 | 0 | 0 | 0 | 1 | 1 |
| 14 | 1 | 1 | 0 | 0 | 0 | 0 | 1 | 1 |
| 15 | 1 | 1 | 0 | 0 | 0 | 0 | 1 | 1 |
| 16 | 1 | 1 | 0 | 0 | 0 | 0 | 1 | 1 |
| 17 | 1 | 1 | 0 | 0 | 0 | 0 | 1 | 1 |
| 18 | 1 | 1 | 0 | 0 | 0 | 0 | 1 | 1 |
| 19 | 1 | 1 | 0 | 0 | 0 | 0 | 1 | 1 |
| 20 | 1 | 1 | 0 | 0 | 0 | 0 | 1 | 1 |
| 1 | 2 | 0 | 1 | 0 | 0 | 0 | 1 | 1 |
| 2 | 2 | 0 | 1 | 0 | 0 | 0 | 1 | 1 |
| 3 | 2 | 0 | 1 | 0 | 0 | 0 | 1 | 1 |
| 4 | 2 | 0 | 1 | 0 | 0 | 0 | 1 | 1 |
| 5 | 2 | 0 | 1 | 0 | 0 | 0 | 1 | 1 |
| 6 | 2 | 0 | 1 | 0 | 0 | 0 | 1 | 1 |
| 7 | 2 | 0 | 1 | 0 | 0 | 0 | 1 | 1 |
| 8 | 2 | 0 | 1 | 0 | 0 | 0 | 1 | 1 |
| 9 | 2 | 0 | 1 | 0 | 0 | 0 | 1 | 1 |
| 10 | 2 | 0 | 1 | 0 | 0 | 0 | 1 | 1 |
| 11 | 2 | 0 | 1 | 0 | 0 | 0 | 1 | 1 |
| 12 | 2 | 0 | 1 | 0 | 0 | 0 | 1 | 1 |
| 13 | 2 | 0 | 1 | 0 | 0 | 0 | 1 | 1 |
| 14 | 2 | 0 | 1 | 0 | 0 | 0 | 1 | 1 |
| 15 | 2 | 0 | 1 | 0 | 0 | 0 | 1 | 1 |
| 16 | 2 | 0 | 1 | 0 | 0 | 0 | 1 | 1 |
| 17 | 2 | 0 | 1 | 0 | 0 | 0 | 1 | 1 |
| 18 | 2 | 0 | 1 | 0 | 0 | 0 | 1 | 1 |
| 19 | 2 | 0 | 1 | 0 | 0 | 0 | 1 | 1 |
| 20 | 2 | 0 | 1 | 0 | 0 | 0 | 1 | 1 |
| 1 | 3 | 0 | 0 | 1 | 0 | 0 | 1 | 1 |
| 2 | 3 | 0 | 0 | 1 | 0 | 0 | 1 | 1 |
| 3 | 3 | 0 | 0 | 1 | 0 | 0 | 1 | 1 |
| 4 | 3 | 0 | 0 | 1 | 0 | 0 | 1 | 1 |
| 5 | 3 | 0 | 0 | 1 | 0 | 0 | 1 | 1 |
| 6 | 3 | 0 | 0 | 1 | 0 | 0 | 1 | 1 |
| 7 | 3 | 0 | 0 | 1 | 0 | 0 | 1 | 1 |
| 8 | 3 | 0 | 0 | 1 | 0 | 0 | 1 | 1 |
| 9 | 3 | 0 | 0 | 1 | 0 | 0 | 1 | 1 |
| 10 | 3 | 0 | 0 | 1 | 0 | 0 | 1 | 1 |
| 11 | 3 | 0 | 0 | 1 | 0 | 0 | 1 | 1 |
| 12 | 3 | 0 | 0 | 1 | 0 | 0 | 1 | 1 |
| 13 | 3 | 0 | 0 | 1 | 0 | 0 | 1 | 1 |
| 14 | 3 | 0 | 0 | 1 | 0 | 0 | 1 | 1 |
| 15 | 3 | 0 | 0 | 1 | 0 | 0 | 1 | 1 |
| 16 | 3 | 0 | 0 | 1 | 0 | 0 | 1 | 1 |
| 17 | 3 | 0 | 0 | 1 | 0 | 0 | 1 | 1 |
| 18 | 3 | 0 | 0 | 1 | 0 | 0 | 1 | 1 |
| 19 | 3 | 0 | 0 | 1 | 0 | 0 | 1 | 1 |
| 20 | 3 | 0 | 0 | 1 | 0 | 0 | 1 | 1 |
| 1 | 4 | 0 | 0 | 0 | 1 | 0 | 1 | 1 |
| 2 | 4 | 0 | 0 | 0 | 1 | 0 | 1 | 1 |
| 3 | 4 | 0 | 0 | 0 | 1 | 0 | 1 | 1 |
| 4 | 4 | 0 | 0 | 0 | 1 | 0 | 1 | 1 |
| 5 | 4 | 0 | 0 | 0 | 1 | 0 | 1 | 1 |
| 6 | 4 | 0 | 0 | 0 | 1 | 0 | 1 | 1 |
| 7 | 4 | 0 | 0 | 0 | 1 | 0 | 1 | 1 |
| 8 | 4 | 0 | 0 | 0 | 1 | 0 | 1 | 1 |
| 9 | 4 | 0 | 0 | 0 | 1 | 0 | 1 | 1 |
| 10 | 4 | 0 | 0 | 0 | 1 | 0 | 1 | 1 |
| 11 | 4 | 0 | 0 | 0 | 1 | 0 | 1 | 1 |
| 12 | 4 | 0 | 0 | 0 | 1 | 0 | 1 | 1 |
| 13 | 4 | 0 | 0 | 0 | 1 | 0 | 1 | 1 |
| 14 | 4 | 0 | 0 | 0 | 1 | 0 | 1 | 1 |
| 15 | 4 | 0 | 0 | 0 | 1 | 0 | 1 | 1 |
| 16 | 4 | 0 | 0 | 0 | 1 | 0 | 1 | 1 |
| 17 | 4 | 0 | 0 | 0 | 1 | 0 | 1 | 1 |
| 18 | 4 | 0 | 0 | 0 | 1 | 0 | 1 | 1 |
| 19 | 4 | 0 | 0 | 0 | 1 | 0 | 1 | 1 |
| 20 | 4 | 0 | 0 | 0 | 1 | 0 | 1 | 1 |
| 1 | 5 | 0 | 0 | 0 | 0 | 1 | 1 | 1 |
| 2 | 5 | 0 | 0 | 0 | 0 | 1 | 1 | 1 |
| 3 | 5 | 0 | 0 | 0 | 0 | 1 | 1 | 1 |
| 4 | 5 | 0 | 0 | 0 | 0 | 1 | 1 | 1 |
| 5 | 5 | 0 | 0 | 0 | 0 | 1 | 1 | 1 |
| 6 | 5 | 0 | 0 | 0 | 0 | 1 | 1 | 1 |
| 7 | 5 | 0 | 0 | 0 | 0 | 1 | 1 | 1 |
| 8 | 5 | 0 | 0 | 0 | 0 | 1 | 1 | 1 |
| 9 | 5 | 0 | 0 | 0 | 0 | 1 | 1 | 1 |
| 10 | 5 | 0 | 0 | 0 | 0 | 1 | 1 | 1 |
| 11 | 5 | 0 | 0 | 0 | 0 | 1 | 1 | 1 |
| 12 | 5 | 0 | 0 | 0 | 0 | 1 | 1 | 1 |
| 13 | 5 | 0 | 0 | 0 | 0 | 1 | 1 | 1 |
| 14 | 5 | 0 | 0 | 0 | 0 | 1 | 1 | 1 |
| 15 | 5 | 0 | 0 | 0 | 0 | 1 | 1 | 1 |
| 16 | 5 | 0 | 0 | 0 | 0 | 1 | 1 | 1 |
| 17 | 5 | 0 | 0 | 0 | 0 | 1 | 1 | 1 |
| 18 | 5 | 0 | 0 | 0 | 0 | 1 | 1 | 1 |
| 19 | 5 | 0 | 0 | 0 | 0 | 1 | 1 | 1 |
| 20 | 5 | 0 | 0 | 0 | 0 | 1 | 1 | 1 |
| 1 | 6 | 1 | 1 | 0 | 0 | 0 | 1 | 2 |
| 2 | 6 | 1 | 1 | 0 | 0 | 0 | 1 | 2 |
| 3 | 6 | 1 | 1 | 0 | 0 | 0 | 1 | 2 |
| 4 | 6 | 1 | 1 | 0 | 0 | 0 | 1 | 2 |
| 5 | 6 | 1 | 1 | 0 | 0 | 0 | 1 | 2 |
| 6 | 6 | 1 | 1 | 0 | 0 | 0 | 1 | 2 |
| 7 | 6 | 1 | 1 | 0 | 0 | 0 | 1 | 2 |
| 8 | 6 | 1 | 1 | 0 | 0 | 0 | 1 | 2 |
| 9 | 6 | 1 | 1 | 0 | 0 | 0 | 1 | 2 |
| 10 | 6 | 1 | 1 | 0 | 0 | 0 | 1 | 2 |
| 11 | 6 | 1 | 1 | 0 | 0 | 0 | 1 | 2 |
| 12 | 6 | 1 | 1 | 0 | 0 | 0 | 1 | 2 |
| 13 | 6 | 1 | 1 | 0 | 0 | 0 | 1 | 2 |
| 14 | 6 | 1 | 1 | 0 | 0 | 0 | 1 | 2 |
| 15 | 6 | 1 | 1 | 0 | 0 | 0 | 1 | 2 |
| 16 | 6 | 1 | 1 | 0 | 0 | 0 | 1 | 2 |
| 17 | 6 | 1 | 1 | 0 | 0 | 0 | 1 | 2 |
| 18 | 6 | 1 | 1 | 0 | 0 | 0 | 1 | 2 |
| 19 | 6 | 1 | 1 | 0 | 0 | 0 | 1 | 2 |
| 20 | 6 | 1 | 1 | 0 | 0 | 0 | 1 | 2 |
| 1 | 7 | 1 | 0 | 1 | 0 | 0 | 1 | 2 |
| 2 | 7 | 1 | 0 | 1 | 0 | 0 | 1 | 2 |
| 3 | 7 | 1 | 0 | 1 | 0 | 0 | 1 | 2 |
| 4 | 7 | 1 | 0 | 1 | 0 | 0 | 1 | 2 |
| 5 | 7 | 1 | 0 | 1 | 0 | 0 | 1 | 2 |
| 6 | 7 | 1 | 0 | 1 | 0 | 0 | 1 | 2 |
| 7 | 7 | 1 | 0 | 1 | 0 | 0 | 1 | 2 |
| 8 | 7 | 1 | 0 | 1 | 0 | 0 | 1 | 2 |
| 9 | 7 | 1 | 0 | 1 | 0 | 0 | 1 | 2 |
| 10 | 7 | 1 | 0 | 1 | 0 | 0 | 1 | 2 |
| 11 | 7 | 1 | 0 | 1 | 0 | 0 | 1 | 2 |
| 12 | 7 | 1 | 0 | 1 | 0 | 0 | 1 | 2 |
| 13 | 7 | 1 | 0 | 1 | 0 | 0 | 1 | 2 |
| 14 | 7 | 1 | 0 | 1 | 0 | 0 | 1 | 2 |
| 15 | 7 | 1 | 0 | 1 | 0 | 0 | 1 | 2 |
| 16 | 7 | 1 | 0 | 1 | 0 | 0 | 1 | 2 |
| 17 | 7 | 1 | 0 | 1 | 0 | 0 | 1 | 2 |
| 18 | 7 | 1 | 0 | 1 | 0 | 0 | 1 | 2 |
| 19 | 7 | 1 | 0 | 1 | 0 | 0 | 1 | 2 |
| 20 | 7 | 1 | 0 | 1 | 0 | 0 | 1 | 2 |
| 1 | 8 | 1 | 0 | 0 | 1 | 0 | 1 | 2 |
| 2 | 8 | 1 | 0 | 0 | 1 | 0 | 1 | 2 |
| 3 | 8 | 1 | 0 | 0 | 1 | 0 | 1 | 2 |
| 4 | 8 | 1 | 0 | 0 | 1 | 0 | 1 | 2 |
| 5 | 8 | 1 | 0 | 0 | 1 | 0 | 1 | 2 |
| 6 | 8 | 1 | 0 | 0 | 1 | 0 | 1 | 2 |
| 7 | 8 | 1 | 0 | 0 | 1 | 0 | 1 | 2 |
| 8 | 8 | 1 | 0 | 0 | 1 | 0 | 1 | 2 |
| 9 | 8 | 1 | 0 | 0 | 1 | 0 | 1 | 2 |
| 10 | 8 | 1 | 0 | 0 | 1 | 0 | 1 | 2 |
| 11 | 8 | 1 | 0 | 0 | 1 | 0 | 1 | 2 |
| 12 | 8 | 1 | 0 | 0 | 1 | 0 | 1 | 2 |
| 13 | 8 | 1 | 0 | 0 | 1 | 0 | 1 | 2 |
| 14 | 8 | 1 | 0 | 0 | 1 | 0 | 1 | 2 |
| 15 | 8 | 1 | 0 | 0 | 1 | 0 | 1 | 2 |
| 16 | 8 | 1 | 0 | 0 | 1 | 0 | 1 | 2 |
| 17 | 8 | 1 | 0 | 0 | 1 | 0 | 1 | 2 |
| 18 | 8 | 1 | 0 | 0 | 1 | 0 | 1 | 2 |
| 19 | 8 | 1 | 0 | 0 | 1 | 0 | 1 | 2 |
| 20 | 8 | 1 | 0 | 0 | 1 | 0 | 1 | 2 |
| 1 | 9 | 1 | 0 | 0 | 0 | 1 | 1 | 2 |
| 2 | 9 | 1 | 0 | 0 | 0 | 1 | 1 | 2 |
| 3 | 9 | 1 | 0 | 0 | 0 | 1 | 1 | 2 |
| 4 | 9 | 1 | 0 | 0 | 0 | 1 | 1 | 2 |
| 5 | 9 | 1 | 0 | 0 | 0 | 1 | 1 | 2 |
| 6 | 9 | 1 | 0 | 0 | 0 | 1 | 1 | 2 |
| 7 | 9 | 1 | 0 | 0 | 0 | 1 | 1 | 2 |
| 8 | 9 | 1 | 0 | 0 | 0 | 1 | 1 | 2 |
| 9 | 9 | 1 | 0 | 0 | 0 | 1 | 1 | 2 |
| 10 | 9 | 1 | 0 | 0 | 0 | 1 | 1 | 2 |
| 11 | 9 | 1 | 0 | 0 | 0 | 1 | 1 | 2 |
| 12 | 9 | 1 | 0 | 0 | 0 | 1 | 1 | 2 |
| 13 | 9 | 1 | 0 | 0 | 0 | 1 | 1 | 2 |
| 14 | 9 | 1 | 0 | 0 | 0 | 1 | 1 | 2 |
| 15 | 9 | 1 | 0 | 0 | 0 | 1 | 1 | 2 |
| 16 | 9 | 1 | 0 | 0 | 0 | 1 | 1 | 2 |
| 17 | 9 | 1 | 0 | 0 | 0 | 1 | 1 | 2 |
| 18 | 9 | 1 | 0 | 0 | 0 | 1 | 1 | 2 |
| 19 | 9 | 1 | 0 | 0 | 0 | 1 | 1 | 2 |
| 20 | 9 | 1 | 0 | 0 | 0 | 1 | 1 | 2 |
| 1 | 10 | 0 | 1 | 1 | 0 | 0 | 1 | 2 |
| 2 | 10 | 0 | 1 | 1 | 0 | 0 | 1 | 2 |
| 3 | 10 | 0 | 1 | 1 | 0 | 0 | 1 | 2 |
| 4 | 10 | 0 | 1 | 1 | 0 | 0 | 1 | 2 |
| 5 | 10 | 0 | 1 | 1 | 0 | 0 | 1 | 2 |
| 6 | 10 | 0 | 1 | 1 | 0 | 0 | 1 | 2 |
| 7 | 10 | 0 | 1 | 1 | 0 | 0 | 1 | 2 |
| 8 | 10 | 0 | 1 | 1 | 0 | 0 | 1 | 2 |
| 9 | 10 | 0 | 1 | 1 | 0 | 0 | 1 | 2 |
| 10 | 10 | 0 | 1 | 1 | 0 | 0 | 1 | 2 |
| 11 | 10 | 0 | 1 | 1 | 0 | 0 | 1 | 2 |
| 12 | 10 | 0 | 1 | 1 | 0 | 0 | 1 | 2 |
| 13 | 10 | 0 | 1 | 1 | 0 | 0 | 1 | 2 |
| 14 | 10 | 0 | 1 | 1 | 0 | 0 | 1 | 2 |
| 15 | 10 | 0 | 1 | 1 | 0 | 0 | 1 | 2 |
| 16 | 10 | 0 | 1 | 1 | 0 | 0 | 1 | 2 |
| 17 | 10 | 0 | 1 | 1 | 0 | 0 | 1 | 2 |
| 18 | 10 | 0 | 1 | 1 | 0 | 0 | 1 | 2 |
| 19 | 10 | 0 | 1 | 1 | 0 | 0 | 1 | 2 |
| 20 | 10 | 0 | 1 | 1 | 0 | 0 | 1 | 2 |
| 1 | 11 | 0 | 1 | 0 | 1 | 0 | 1 | 2 |
| 2 | 11 | 0 | 1 | 0 | 1 | 0 | 1 | 2 |
| 3 | 11 | 0 | 1 | 0 | 1 | 0 | 1 | 2 |
| 4 | 11 | 0 | 1 | 0 | 1 | 0 | 1 | 2 |
| 5 | 11 | 0 | 1 | 0 | 1 | 0 | 1 | 2 |
| 6 | 11 | 0 | 1 | 0 | 1 | 0 | 1 | 2 |
| 7 | 11 | 0 | 1 | 0 | 1 | 0 | 1 | 2 |
| 8 | 11 | 0 | 1 | 0 | 1 | 0 | 1 | 2 |
| 9 | 11 | 0 | 1 | 0 | 1 | 0 | 1 | 2 |
| 10 | 11 | 0 | 1 | 0 | 1 | 0 | 1 | 2 |
| 11 | 11 | 0 | 1 | 0 | 1 | 0 | 1 | 2 |
| 12 | 11 | 0 | 1 | 0 | 1 | 0 | 1 | 2 |
| 13 | 11 | 0 | 1 | 0 | 1 | 0 | 1 | 2 |
| 14 | 11 | 0 | 1 | 0 | 1 | 0 | 1 | 2 |
| 15 | 11 | 0 | 1 | 0 | 1 | 0 | 1 | 2 |
| 16 | 11 | 0 | 1 | 0 | 1 | 0 | 1 | 2 |
| 17 | 11 | 0 | 1 | 0 | 1 | 0 | 1 | 2 |
| 18 | 11 | 0 | 1 | 0 | 1 | 0 | 1 | 2 |
| 19 | 11 | 0 | 1 | 0 | 1 | 0 | 1 | 2 |
| 20 | 11 | 0 | 1 | 0 | 1 | 0 | 1 | 2 |
| 1 | 12 | 0 | 1 | 0 | 0 | 1 | 1 | 2 |
| 2 | 12 | 0 | 1 | 0 | 0 | 1 | 1 | 2 |
| 3 | 12 | 0 | 1 | 0 | 0 | 1 | 1 | 2 |
| 4 | 12 | 0 | 1 | 0 | 0 | 1 | 1 | 2 |
| 5 | 12 | 0 | 1 | 0 | 0 | 1 | 1 | 2 |
| 6 | 12 | 0 | 1 | 0 | 0 | 1 | 1 | 2 |
| 7 | 12 | 0 | 1 | 0 | 0 | 1 | 1 | 2 |
| 8 | 12 | 0 | 1 | 0 | 0 | 1 | 1 | 2 |
| 9 | 12 | 0 | 1 | 0 | 0 | 1 | 1 | 2 |
| 10 | 12 | 0 | 1 | 0 | 0 | 1 | 1 | 2 |
| 11 | 12 | 0 | 1 | 0 | 0 | 1 | 1 | 2 |
| 12 | 12 | 0 | 1 | 0 | 0 | 1 | 1 | 2 |
| 13 | 12 | 0 | 1 | 0 | 0 | 1 | 1 | 2 |
| 14 | 12 | 0 | 1 | 0 | 0 | 1 | 1 | 2 |
| 15 | 12 | 0 | 1 | 0 | 0 | 1 | 1 | 2 |
| 16 | 12 | 0 | 1 | 0 | 0 | 1 | 1 | 2 |
| 17 | 12 | 0 | 1 | 0 | 0 | 1 | 1 | 2 |
| 18 | 12 | 0 | 1 | 0 | 0 | 1 | 1 | 2 |
| 19 | 12 | 0 | 1 | 0 | 0 | 1 | 1 | 2 |
| 20 | 12 | 0 | 1 | 0 | 0 | 1 | 1 | 2 |
| 1 | 13 | 0 | 0 | 1 | 1 | 0 | 1 | 2 |
| 2 | 13 | 0 | 0 | 1 | 1 | 0 | 1 | 2 |
| 3 | 13 | 0 | 0 | 1 | 1 | 0 | 1 | 2 |
| 4 | 13 | 0 | 0 | 1 | 1 | 0 | 1 | 2 |
| 5 | 13 | 0 | 0 | 1 | 1 | 0 | 1 | 2 |
| 6 | 13 | 0 | 0 | 1 | 1 | 0 | 1 | 2 |
| 7 | 13 | 0 | 0 | 1 | 1 | 0 | 1 | 2 |
| 8 | 13 | 0 | 0 | 1 | 1 | 0 | 1 | 2 |
| 9 | 13 | 0 | 0 | 1 | 1 | 0 | 1 | 2 |
| 10 | 13 | 0 | 0 | 1 | 1 | 0 | 1 | 2 |
| 11 | 13 | 0 | 0 | 1 | 1 | 0 | 1 | 2 |
| 12 | 13 | 0 | 0 | 1 | 1 | 0 | 1 | 2 |
| 13 | 13 | 0 | 0 | 1 | 1 | 0 | 1 | 2 |
| 14 | 13 | 0 | 0 | 1 | 1 | 0 | 1 | 2 |
| 15 | 13 | 0 | 0 | 1 | 1 | 0 | 1 | 2 |
| 16 | 13 | 0 | 0 | 1 | 1 | 0 | 1 | 2 |
| 17 | 13 | 0 | 0 | 1 | 1 | 0 | 1 | 2 |
| 18 | 13 | 0 | 0 | 1 | 1 | 0 | 1 | 2 |
| 19 | 13 | 0 | 0 | 1 | 1 | 0 | 1 | 2 |
| 20 | 13 | 0 | 0 | 1 | 1 | 0 | 1 | 2 |
| 1 | 14 | 0 | 0 | 1 | 0 | 1 | 1 | 2 |
| 2 | 14 | 0 | 0 | 1 | 0 | 1 | 1 | 2 |
| 3 | 14 | 0 | 0 | 1 | 0 | 1 | 1 | 2 |
| 4 | 14 | 0 | 0 | 1 | 0 | 1 | 1 | 2 |
| 5 | 14 | 0 | 0 | 1 | 0 | 1 | 1 | 2 |
| 6 | 14 | 0 | 0 | 1 | 0 | 1 | 1 | 2 |
| 7 | 14 | 0 | 0 | 1 | 0 | 1 | 1 | 2 |
| 8 | 14 | 0 | 0 | 1 | 0 | 1 | 1 | 2 |
| 9 | 14 | 0 | 0 | 1 | 0 | 1 | 1 | 2 |
| 10 | 14 | 0 | 0 | 1 | 0 | 1 | 1 | 2 |
| 11 | 14 | 0 | 0 | 1 | 0 | 1 | 1 | 2 |
| 12 | 14 | 0 | 0 | 1 | 0 | 1 | 1 | 2 |
| 13 | 14 | 0 | 0 | 1 | 0 | 1 | 1 | 2 |
| 14 | 14 | 0 | 0 | 1 | 0 | 1 | 1 | 2 |
| 15 | 14 | 0 | 0 | 1 | 0 | 1 | 1 | 2 |
| 16 | 14 | 0 | 0 | 1 | 0 | 1 | 1 | 2 |
| 17 | 14 | 0 | 0 | 1 | 0 | 1 | 1 | 2 |
| 18 | 14 | 0 | 0 | 1 | 0 | 1 | 1 | 2 |
| 19 | 14 | 0 | 0 | 1 | 0 | 1 | 1 | 2 |
| 20 | 14 | 0 | 0 | 1 | 0 | 1 | 1 | 2 |
| 1 | 15 | 0 | 0 | 0 | 1 | 1 | 1 | 2 |
| 2 | 15 | 0 | 0 | 0 | 1 | 1 | 1 | 2 |
| 3 | 15 | 0 | 0 | 0 | 1 | 1 | 1 | 2 |
| 4 | 15 | 0 | 0 | 0 | 1 | 1 | 1 | 2 |
| 5 | 15 | 0 | 0 | 0 | 1 | 1 | 1 | 2 |
| 6 | 15 | 0 | 0 | 0 | 1 | 1 | 1 | 2 |
| 7 | 15 | 0 | 0 | 0 | 1 | 1 | 1 | 2 |
| 8 | 15 | 0 | 0 | 0 | 1 | 1 | 1 | 2 |
| 9 | 15 | 0 | 0 | 0 | 1 | 1 | 1 | 2 |
| 10 | 15 | 0 | 0 | 0 | 1 | 1 | 1 | 2 |
| 11 | 15 | 0 | 0 | 0 | 1 | 1 | 1 | 2 |
| 12 | 15 | 0 | 0 | 0 | 1 | 1 | 1 | 2 |
| 13 | 15 | 0 | 0 | 0 | 1 | 1 | 1 | 2 |
| 14 | 15 | 0 | 0 | 0 | 1 | 1 | 1 | 2 |
| 15 | 15 | 0 | 0 | 0 | 1 | 1 | 1 | 2 |
| 16 | 15 | 0 | 0 | 0 | 1 | 1 | 1 | 2 |
| 17 | 15 | 0 | 0 | 0 | 1 | 1 | 1 | 2 |
| 18 | 15 | 0 | 0 | 0 | 1 | 1 | 1 | 2 |
| 19 | 15 | 0 | 0 | 0 | 1 | 1 | 1 | 2 |
| 20 | 15 | 0 | 0 | 0 | 1 | 1 | 1 | 2 |
| 1 | 16 | 1 | 1 | 1 | 0 | 0 | 1 | 3 |
| 2 | 16 | 1 | 1 | 1 | 0 | 0 | 1 | 3 |
| 3 | 16 | 1 | 1 | 1 | 0 | 0 | 1 | 3 |
| 4 | 16 | 1 | 1 | 1 | 0 | 0 | 1 | 3 |
| 5 | 16 | 1 | 1 | 1 | 0 | 0 | 1 | 3 |
| 6 | 16 | 1 | 1 | 1 | 0 | 0 | 1 | 3 |
| 7 | 16 | 1 | 1 | 1 | 0 | 0 | 1 | 3 |
| 8 | 16 | 1 | 1 | 1 | 0 | 0 | 1 | 3 |
| 9 | 16 | 1 | 1 | 1 | 0 | 0 | 1 | 3 |
| 10 | 16 | 1 | 1 | 1 | 0 | 0 | 1 | 3 |
| 11 | 16 | 1 | 1 | 1 | 0 | 0 | 1 | 3 |
| 12 | 16 | 1 | 1 | 1 | 0 | 0 | 1 | 3 |
| 13 | 16 | 1 | 1 | 1 | 0 | 0 | 1 | 3 |
| 14 | 16 | 1 | 1 | 1 | 0 | 0 | 1 | 3 |
| 15 | 16 | 1 | 1 | 1 | 0 | 0 | 1 | 3 |
| 16 | 16 | 1 | 1 | 1 | 0 | 0 | 1 | 3 |
| 17 | 16 | 1 | 1 | 1 | 0 | 0 | 1 | 3 |
| 18 | 16 | 1 | 1 | 1 | 0 | 0 | 1 | 3 |
| 19 | 16 | 1 | 1 | 1 | 0 | 0 | 1 | 3 |
| 20 | 16 | 1 | 1 | 1 | 0 | 0 | 1 | 3 |
| 1 | 17 | 1 | 1 | 0 | 1 | 0 | 1 | 3 |
| 2 | 17 | 1 | 1 | 0 | 1 | 0 | 1 | 3 |
| 3 | 17 | 1 | 1 | 0 | 1 | 0 | 1 | 3 |
| 4 | 17 | 1 | 1 | 0 | 1 | 0 | 1 | 3 |
| 5 | 17 | 1 | 1 | 0 | 1 | 0 | 1 | 3 |
| 6 | 17 | 1 | 1 | 0 | 1 | 0 | 1 | 3 |
| 7 | 17 | 1 | 1 | 0 | 1 | 0 | 1 | 3 |
| 8 | 17 | 1 | 1 | 0 | 1 | 0 | 1 | 3 |
| 9 | 17 | 1 | 1 | 0 | 1 | 0 | 1 | 3 |
| 10 | 17 | 1 | 1 | 0 | 1 | 0 | 1 | 3 |
| 11 | 17 | 1 | 1 | 0 | 1 | 0 | 1 | 3 |
| 12 | 17 | 1 | 1 | 0 | 1 | 0 | 1 | 3 |
| 13 | 17 | 1 | 1 | 0 | 1 | 0 | 1 | 3 |
| 14 | 17 | 1 | 1 | 0 | 1 | 0 | 1 | 3 |
| 15 | 17 | 1 | 1 | 0 | 1 | 0 | 1 | 3 |
| 16 | 17 | 1 | 1 | 0 | 1 | 0 | 1 | 3 |
| 17 | 17 | 1 | 1 | 0 | 1 | 0 | 1 | 3 |
| 18 | 17 | 1 | 1 | 0 | 1 | 0 | 1 | 3 |
| 19 | 17 | 1 | 1 | 0 | 1 | 0 | 1 | 3 |
| 20 | 17 | 1 | 1 | 0 | 1 | 0 | 1 | 3 |
| 1 | 18 | 1 | 1 | 0 | 0 | 1 | 1 | 3 |
| 2 | 18 | 1 | 1 | 0 | 0 | 1 | 1 | 3 |
| 3 | 18 | 1 | 1 | 0 | 0 | 1 | 1 | 3 |
| 4 | 18 | 1 | 1 | 0 | 0 | 1 | 1 | 3 |
| 5 | 18 | 1 | 1 | 0 | 0 | 1 | 1 | 3 |
| 6 | 18 | 1 | 1 | 0 | 0 | 1 | 1 | 3 |
| 7 | 18 | 1 | 1 | 0 | 0 | 1 | 1 | 3 |
| 8 | 18 | 1 | 1 | 0 | 0 | 1 | 1 | 3 |
| 9 | 18 | 1 | 1 | 0 | 0 | 1 | 1 | 3 |
| 10 | 18 | 1 | 1 | 0 | 0 | 1 | 1 | 3 |
| 11 | 18 | 1 | 1 | 0 | 0 | 1 | 1 | 3 |
| 12 | 18 | 1 | 1 | 0 | 0 | 1 | 1 | 3 |
| 13 | 18 | 1 | 1 | 0 | 0 | 1 | 1 | 3 |
| 14 | 18 | 1 | 1 | 0 | 0 | 1 | 1 | 3 |
| 15 | 18 | 1 | 1 | 0 | 0 | 1 | 1 | 3 |
| 16 | 18 | 1 | 1 | 0 | 0 | 1 | 1 | 3 |
| 17 | 18 | 1 | 1 | 0 | 0 | 1 | 1 | 3 |
| 18 | 18 | 1 | 1 | 0 | 0 | 1 | 1 | 3 |
| 19 | 18 | 1 | 1 | 0 | 0 | 1 | 1 | 3 |
| 20 | 18 | 1 | 1 | 0 | 0 | 1 | 1 | 3 |
| 1 | 19 | 1 | 0 | 1 | 1 | 0 | 1 | 3 |
| 2 | 19 | 1 | 0 | 1 | 1 | 0 | 1 | 3 |
| 3 | 19 | 1 | 0 | 1 | 1 | 0 | 1 | 3 |
| 4 | 19 | 1 | 0 | 1 | 1 | 0 | 1 | 3 |
| 5 | 19 | 1 | 0 | 1 | 1 | 0 | 1 | 3 |
| 6 | 19 | 1 | 0 | 1 | 1 | 0 | 1 | 3 |
| 7 | 19 | 1 | 0 | 1 | 1 | 0 | 1 | 3 |
| 8 | 19 | 1 | 0 | 1 | 1 | 0 | 1 | 3 |
| 9 | 19 | 1 | 0 | 1 | 1 | 0 | 1 | 3 |
| 10 | 19 | 1 | 0 | 1 | 1 | 0 | 1 | 3 |
| 11 | 19 | 1 | 0 | 1 | 1 | 0 | 1 | 3 |
| 12 | 19 | 1 | 0 | 1 | 1 | 0 | 1 | 3 |
| 13 | 19 | 1 | 0 | 1 | 1 | 0 | 1 | 3 |
| 14 | 19 | 1 | 0 | 1 | 1 | 0 | 1 | 3 |
| 15 | 19 | 1 | 0 | 1 | 1 | 0 | 1 | 3 |
| 16 | 19 | 1 | 0 | 1 | 1 | 0 | 1 | 3 |
| 17 | 19 | 1 | 0 | 1 | 1 | 0 | 1 | 3 |
| 18 | 19 | 1 | 0 | 1 | 1 | 0 | 1 | 3 |
| 19 | 19 | 1 | 0 | 1 | 1 | 0 | 1 | 3 |
| 20 | 19 | 1 | 0 | 1 | 1 | 0 | 1 | 3 |
| 1 | 20 | 1 | 0 | 1 | 0 | 1 | 1 | 3 |
| 2 | 20 | 1 | 0 | 1 | 0 | 1 | 1 | 3 |
| 3 | 20 | 1 | 0 | 1 | 0 | 1 | 1 | 3 |
| 4 | 20 | 1 | 0 | 1 | 0 | 1 | 1 | 3 |
| 5 | 20 | 1 | 0 | 1 | 0 | 1 | 1 | 3 |
| 6 | 20 | 1 | 0 | 1 | 0 | 1 | 1 | 3 |
| 7 | 20 | 1 | 0 | 1 | 0 | 1 | 1 | 3 |
| 8 | 20 | 1 | 0 | 1 | 0 | 1 | 1 | 3 |
| 9 | 20 | 1 | 0 | 1 | 0 | 1 | 1 | 3 |
| 10 | 20 | 1 | 0 | 1 | 0 | 1 | 1 | 3 |
| 11 | 20 | 1 | 0 | 1 | 0 | 1 | 1 | 3 |
| 12 | 20 | 1 | 0 | 1 | 0 | 1 | 1 | 3 |
| 13 | 20 | 1 | 0 | 1 | 0 | 1 | 1 | 3 |
| 14 | 20 | 1 | 0 | 1 | 0 | 1 | 1 | 3 |
| 15 | 20 | 1 | 0 | 1 | 0 | 1 | 1 | 3 |
| 16 | 20 | 1 | 0 | 1 | 0 | 1 | 1 | 3 |
| 17 | 20 | 1 | 0 | 1 | 0 | 1 | 1 | 3 |
| 18 | 20 | 1 | 0 | 1 | 0 | 1 | 1 | 3 |
| 19 | 20 | 1 | 0 | 1 | 0 | 1 | 1 | 3 |
| 20 | 20 | 1 | 0 | 1 | 0 | 1 | 1 | 3 |
| 1 | 21 | 1 | 0 | 0 | 1 | 1 | 1 | 3 |
| 2 | 21 | 1 | 0 | 0 | 1 | 1 | 1 | 3 |
| 3 | 21 | 1 | 0 | 0 | 1 | 1 | 1 | 3 |
| 4 | 21 | 1 | 0 | 0 | 1 | 1 | 1 | 3 |
| 5 | 21 | 1 | 0 | 0 | 1 | 1 | 1 | 3 |
| 6 | 21 | 1 | 0 | 0 | 1 | 1 | 1 | 3 |
| 7 | 21 | 1 | 0 | 0 | 1 | 1 | 1 | 3 |
| 8 | 21 | 1 | 0 | 0 | 1 | 1 | 1 | 3 |
| 9 | 21 | 1 | 0 | 0 | 1 | 1 | 1 | 3 |
| 10 | 21 | 1 | 0 | 0 | 1 | 1 | 1 | 3 |
| 11 | 21 | 1 | 0 | 0 | 1 | 1 | 1 | 3 |
| 12 | 21 | 1 | 0 | 0 | 1 | 1 | 1 | 3 |
| 13 | 21 | 1 | 0 | 0 | 1 | 1 | 1 | 3 |
| 14 | 21 | 1 | 0 | 0 | 1 | 1 | 1 | 3 |
| 15 | 21 | 1 | 0 | 0 | 1 | 1 | 1 | 3 |
| 16 | 21 | 1 | 0 | 0 | 1 | 1 | 1 | 3 |
| 17 | 21 | 1 | 0 | 0 | 1 | 1 | 1 | 3 |
| 18 | 21 | 1 | 0 | 0 | 1 | 1 | 1 | 3 |
| 19 | 21 | 1 | 0 | 0 | 1 | 1 | 1 | 3 |
| 20 | 21 | 1 | 0 | 0 | 1 | 1 | 1 | 3 |
| 1 | 22 | 0 | 1 | 1 | 1 | 0 | 1 | 3 |
| 2 | 22 | 0 | 1 | 1 | 1 | 0 | 1 | 3 |
| 3 | 22 | 0 | 1 | 1 | 1 | 0 | 1 | 3 |
| 4 | 22 | 0 | 1 | 1 | 1 | 0 | 1 | 3 |
| 5 | 22 | 0 | 1 | 1 | 1 | 0 | 1 | 3 |
| 6 | 22 | 0 | 1 | 1 | 1 | 0 | 1 | 3 |
| 7 | 22 | 0 | 1 | 1 | 1 | 0 | 1 | 3 |
| 8 | 22 | 0 | 1 | 1 | 1 | 0 | 1 | 3 |
| 9 | 22 | 0 | 1 | 1 | 1 | 0 | 1 | 3 |
| 10 | 22 | 0 | 1 | 1 | 1 | 0 | 1 | 3 |
| 11 | 22 | 0 | 1 | 1 | 1 | 0 | 1 | 3 |
| 12 | 22 | 0 | 1 | 1 | 1 | 0 | 1 | 3 |
| 13 | 22 | 0 | 1 | 1 | 1 | 0 | 1 | 3 |
| 14 | 22 | 0 | 1 | 1 | 1 | 0 | 1 | 3 |
| 15 | 22 | 0 | 1 | 1 | 1 | 0 | 1 | 3 |
| 16 | 22 | 0 | 1 | 1 | 1 | 0 | 1 | 3 |
| 17 | 22 | 0 | 1 | 1 | 1 | 0 | 1 | 3 |
| 18 | 22 | 0 | 1 | 1 | 1 | 0 | 1 | 3 |
| 19 | 22 | 0 | 1 | 1 | 1 | 0 | 1 | 3 |
| 20 | 22 | 0 | 1 | 1 | 1 | 0 | 1 | 3 |
| 1 | 23 | 0 | 1 | 1 | 0 | 1 | 1 | 3 |
| 2 | 23 | 0 | 1 | 1 | 0 | 1 | 1 | 3 |
| 3 | 23 | 0 | 1 | 1 | 0 | 1 | 1 | 3 |
| 4 | 23 | 0 | 1 | 1 | 0 | 1 | 1 | 3 |
| 5 | 23 | 0 | 1 | 1 | 0 | 1 | 1 | 3 |
| 6 | 23 | 0 | 1 | 1 | 0 | 1 | 1 | 3 |
| 7 | 23 | 0 | 1 | 1 | 0 | 1 | 1 | 3 |
| 8 | 23 | 0 | 1 | 1 | 0 | 1 | 1 | 3 |
| 9 | 23 | 0 | 1 | 1 | 0 | 1 | 1 | 3 |
| 10 | 23 | 0 | 1 | 1 | 0 | 1 | 1 | 3 |
| 11 | 23 | 0 | 1 | 1 | 0 | 1 | 1 | 3 |
| 12 | 23 | 0 | 1 | 1 | 0 | 1 | 1 | 3 |
| 13 | 23 | 0 | 1 | 1 | 0 | 1 | 1 | 3 |
| 14 | 23 | 0 | 1 | 1 | 0 | 1 | 1 | 3 |
| 15 | 23 | 0 | 1 | 1 | 0 | 1 | 1 | 3 |
| 16 | 23 | 0 | 1 | 1 | 0 | 1 | 1 | 3 |
| 17 | 23 | 0 | 1 | 1 | 0 | 1 | 1 | 3 |
| 18 | 23 | 0 | 1 | 1 | 0 | 1 | 1 | 3 |
| 19 | 23 | 0 | 1 | 1 | 0 | 1 | 1 | 3 |
| 20 | 23 | 0 | 1 | 1 | 0 | 1 | 1 | 3 |
| 1 | 24 | 0 | 1 | 0 | 1 | 1 | 1 | 3 |
| 2 | 24 | 0 | 1 | 0 | 1 | 1 | 1 | 3 |
| 3 | 24 | 0 | 1 | 0 | 1 | 1 | 1 | 3 |
| 4 | 24 | 0 | 1 | 0 | 1 | 1 | 1 | 3 |
| 5 | 24 | 0 | 1 | 0 | 1 | 1 | 1 | 3 |
| 6 | 24 | 0 | 1 | 0 | 1 | 1 | 1 | 3 |
| 7 | 24 | 0 | 1 | 0 | 1 | 1 | 1 | 3 |
| 8 | 24 | 0 | 1 | 0 | 1 | 1 | 1 | 3 |
| 9 | 24 | 0 | 1 | 0 | 1 | 1 | 1 | 3 |
| 10 | 24 | 0 | 1 | 0 | 1 | 1 | 1 | 3 |
| 11 | 24 | 0 | 1 | 0 | 1 | 1 | 1 | 3 |
| 12 | 24 | 0 | 1 | 0 | 1 | 1 | 1 | 3 |
| 13 | 24 | 0 | 1 | 0 | 1 | 1 | 1 | 3 |
| 14 | 24 | 0 | 1 | 0 | 1 | 1 | 1 | 3 |
| 15 | 24 | 0 | 1 | 0 | 1 | 1 | 1 | 3 |
| 16 | 24 | 0 | 1 | 0 | 1 | 1 | 1 | 3 |
| 17 | 24 | 0 | 1 | 0 | 1 | 1 | 1 | 3 |
| 18 | 24 | 0 | 1 | 0 | 1 | 1 | 1 | 3 |
| 19 | 24 | 0 | 1 | 0 | 1 | 1 | 1 | 3 |
| 20 | 24 | 0 | 1 | 0 | 1 | 1 | 1 | 3 |
| 1 | 25 | 0 | 0 | 1 | 1 | 1 | 1 | 3 |
| 2 | 25 | 0 | 0 | 1 | 1 | 1 | 1 | 3 |
| 3 | 25 | 0 | 0 | 1 | 1 | 1 | 1 | 3 |
| 4 | 25 | 0 | 0 | 1 | 1 | 1 | 1 | 3 |
| 5 | 25 | 0 | 0 | 1 | 1 | 1 | 1 | 3 |
| 6 | 25 | 0 | 0 | 1 | 1 | 1 | 1 | 3 |
| 7 | 25 | 0 | 0 | 1 | 1 | 1 | 1 | 3 |
| 8 | 25 | 0 | 0 | 1 | 1 | 1 | 1 | 3 |
| 9 | 25 | 0 | 0 | 1 | 1 | 1 | 1 | 3 |
| 10 | 25 | 0 | 0 | 1 | 1 | 1 | 1 | 3 |
| 11 | 25 | 0 | 0 | 1 | 1 | 1 | 1 | 3 |
| 12 | 25 | 0 | 0 | 1 | 1 | 1 | 1 | 3 |
| 13 | 25 | 0 | 0 | 1 | 1 | 1 | 1 | 3 |
| 14 | 25 | 0 | 0 | 1 | 1 | 1 | 1 | 3 |
| 15 | 25 | 0 | 0 | 1 | 1 | 1 | 1 | 3 |
| 16 | 25 | 0 | 0 | 1 | 1 | 1 | 1 | 3 |
| 17 | 25 | 0 | 0 | 1 | 1 | 1 | 1 | 3 |
| 18 | 25 | 0 | 0 | 1 | 1 | 1 | 1 | 3 |
| 19 | 25 | 0 | 0 | 1 | 1 | 1 | 1 | 3 |
| 20 | 25 | 0 | 0 | 1 | 1 | 1 | 1 | 3 |
| 1 | 26 | 1 | 1 | 1 | 1 | 0 | 1 | 4 |
| 2 | 26 | 1 | 1 | 1 | 1 | 0 | 1 | 4 |
| 3 | 26 | 1 | 1 | 1 | 1 | 0 | 1 | 4 |
| 4 | 26 | 1 | 1 | 1 | 1 | 0 | 1 | 4 |
| 5 | 26 | 1 | 1 | 1 | 1 | 0 | 1 | 4 |
| 6 | 26 | 1 | 1 | 1 | 1 | 0 | 1 | 4 |
| 7 | 26 | 1 | 1 | 1 | 1 | 0 | 1 | 4 |
| 8 | 26 | 1 | 1 | 1 | 1 | 0 | 1 | 4 |
| 9 | 26 | 1 | 1 | 1 | 1 | 0 | 1 | 4 |
| 10 | 26 | 1 | 1 | 1 | 1 | 0 | 1 | 4 |
| 11 | 26 | 1 | 1 | 1 | 1 | 0 | 1 | 4 |
| 12 | 26 | 1 | 1 | 1 | 1 | 0 | 1 | 4 |
| 13 | 26 | 1 | 1 | 1 | 1 | 0 | 1 | 4 |
| 14 | 26 | 1 | 1 | 1 | 1 | 0 | 1 | 4 |
| 15 | 26 | 1 | 1 | 1 | 1 | 0 | 1 | 4 |
| 16 | 26 | 1 | 1 | 1 | 1 | 0 | 1 | 4 |
| 17 | 26 | 1 | 1 | 1 | 1 | 0 | 1 | 4 |
| 18 | 26 | 1 | 1 | 1 | 1 | 0 | 1 | 4 |
| 19 | 26 | 1 | 1 | 1 | 1 | 0 | 1 | 4 |
| 20 | 26 | 1 | 1 | 1 | 1 | 0 | 1 | 4 |
| 1 | 27 | 1 | 1 | 1 | 0 | 1 | 1 | 4 |
| 2 | 27 | 1 | 1 | 1 | 0 | 1 | 1 | 4 |
| 3 | 27 | 1 | 1 | 1 | 0 | 1 | 1 | 4 |
| 4 | 27 | 1 | 1 | 1 | 0 | 1 | 1 | 4 |
| 5 | 27 | 1 | 1 | 1 | 0 | 1 | 1 | 4 |
| 6 | 27 | 1 | 1 | 1 | 0 | 1 | 1 | 4 |
| 7 | 27 | 1 | 1 | 1 | 0 | 1 | 1 | 4 |
| 8 | 27 | 1 | 1 | 1 | 0 | 1 | 1 | 4 |
| 9 | 27 | 1 | 1 | 1 | 0 | 1 | 1 | 4 |
| 10 | 27 | 1 | 1 | 1 | 0 | 1 | 1 | 4 |
| 11 | 27 | 1 | 1 | 1 | 0 | 1 | 1 | 4 |
| 12 | 27 | 1 | 1 | 1 | 0 | 1 | 1 | 4 |
| 13 | 27 | 1 | 1 | 1 | 0 | 1 | 1 | 4 |
| 14 | 27 | 1 | 1 | 1 | 0 | 1 | 1 | 4 |
| 15 | 27 | 1 | 1 | 1 | 0 | 1 | 1 | 4 |
| 16 | 27 | 1 | 1 | 1 | 0 | 1 | 1 | 4 |
| 17 | 27 | 1 | 1 | 1 | 0 | 1 | 1 | 4 |
| 18 | 27 | 1 | 1 | 1 | 0 | 1 | 1 | 4 |
| 19 | 27 | 1 | 1 | 1 | 0 | 1 | 1 | 4 |
| 20 | 27 | 1 | 1 | 1 | 0 | 1 | 1 | 4 |
| 1 | 28 | 1 | 1 | 0 | 1 | 1 | 1 | 4 |
| 2 | 28 | 1 | 1 | 0 | 1 | 1 | 1 | 4 |
| 3 | 28 | 1 | 1 | 0 | 1 | 1 | 1 | 4 |
| 4 | 28 | 1 | 1 | 0 | 1 | 1 | 1 | 4 |
| 5 | 28 | 1 | 1 | 0 | 1 | 1 | 1 | 4 |
| 6 | 28 | 1 | 1 | 0 | 1 | 1 | 1 | 4 |
| 7 | 28 | 1 | 1 | 0 | 1 | 1 | 1 | 4 |
| 8 | 28 | 1 | 1 | 0 | 1 | 1 | 1 | 4 |
| 9 | 28 | 1 | 1 | 0 | 1 | 1 | 1 | 4 |
| 10 | 28 | 1 | 1 | 0 | 1 | 1 | 1 | 4 |
| 11 | 28 | 1 | 1 | 0 | 1 | 1 | 1 | 4 |
| 12 | 28 | 1 | 1 | 0 | 1 | 1 | 1 | 4 |
| 13 | 28 | 1 | 1 | 0 | 1 | 1 | 1 | 4 |
| 14 | 28 | 1 | 1 | 0 | 1 | 1 | 1 | 4 |
| 15 | 28 | 1 | 1 | 0 | 1 | 1 | 1 | 4 |
| 16 | 28 | 1 | 1 | 0 | 1 | 1 | 1 | 4 |
| 17 | 28 | 1 | 1 | 0 | 1 | 1 | 1 | 4 |
| 18 | 28 | 1 | 1 | 0 | 1 | 1 | 1 | 4 |
| 19 | 28 | 1 | 1 | 0 | 1 | 1 | 1 | 4 |
| 20 | 28 | 1 | 1 | 0 | 1 | 1 | 1 | 4 |
| 1 | 29 | 1 | 0 | 1 | 1 | 1 | 1 | 4 |
| 2 | 29 | 1 | 0 | 1 | 1 | 1 | 1 | 4 |
| 3 | 29 | 1 | 0 | 1 | 1 | 1 | 1 | 4 |
| 4 | 29 | 1 | 0 | 1 | 1 | 1 | 1 | 4 |
| 5 | 29 | 1 | 0 | 1 | 1 | 1 | 1 | 4 |
| 6 | 29 | 1 | 0 | 1 | 1 | 1 | 1 | 4 |
| 7 | 29 | 1 | 0 | 1 | 1 | 1 | 1 | 4 |
| 8 | 29 | 1 | 0 | 1 | 1 | 1 | 1 | 4 |
| 9 | 29 | 1 | 0 | 1 | 1 | 1 | 1 | 4 |
| 10 | 29 | 1 | 0 | 1 | 1 | 1 | 1 | 4 |
| 11 | 29 | 1 | 0 | 1 | 1 | 1 | 1 | 4 |
| 12 | 29 | 1 | 0 | 1 | 1 | 1 | 1 | 4 |
| 13 | 29 | 1 | 0 | 1 | 1 | 1 | 1 | 4 |
| 14 | 29 | 1 | 0 | 1 | 1 | 1 | 1 | 4 |
| 15 | 29 | 1 | 0 | 1 | 1 | 1 | 1 | 4 |
| 16 | 29 | 1 | 0 | 1 | 1 | 1 | 1 | 4 |
| 17 | 29 | 1 | 0 | 1 | 1 | 1 | 1 | 4 |
| 18 | 29 | 1 | 0 | 1 | 1 | 1 | 1 | 4 |
| 19 | 29 | 1 | 0 | 1 | 1 | 1 | 1 | 4 |
| 20 | 29 | 1 | 0 | 1 | 1 | 1 | 1 | 4 |
| 1 | 30 | 0 | 1 | 1 | 1 | 1 | 1 | 4 |
| 2 | 30 | 0 | 1 | 1 | 1 | 1 | 1 | 4 |
| 3 | 30 | 0 | 1 | 1 | 1 | 1 | 1 | 4 |
| 4 | 30 | 0 | 1 | 1 | 1 | 1 | 1 | 4 |
| 5 | 30 | 0 | 1 | 1 | 1 | 1 | 1 | 4 |
| 6 | 30 | 0 | 1 | 1 | 1 | 1 | 1 | 4 |
| 7 | 30 | 0 | 1 | 1 | 1 | 1 | 1 | 4 |
| 8 | 30 | 0 | 1 | 1 | 1 | 1 | 1 | 4 |
| 9 | 30 | 0 | 1 | 1 | 1 | 1 | 1 | 4 |
| 10 | 30 | 0 | 1 | 1 | 1 | 1 | 1 | 4 |
| 11 | 30 | 0 | 1 | 1 | 1 | 1 | 1 | 4 |
| 12 | 30 | 0 | 1 | 1 | 1 | 1 | 1 | 4 |
| 13 | 30 | 0 | 1 | 1 | 1 | 1 | 1 | 4 |
| 14 | 30 | 0 | 1 | 1 | 1 | 1 | 1 | 4 |
| 15 | 30 | 0 | 1 | 1 | 1 | 1 | 1 | 4 |
| 16 | 30 | 0 | 1 | 1 | 1 | 1 | 1 | 4 |
| 17 | 30 | 0 | 1 | 1 | 1 | 1 | 1 | 4 |
| 18 | 30 | 0 | 1 | 1 | 1 | 1 | 1 | 4 |
| 19 | 30 | 0 | 1 | 1 | 1 | 1 | 1 | 4 |
| 20 | 30 | 0 | 1 | 1 | 1 | 1 | 1 | 4 |
| 1 | 31 | 1 | 1 | 1 | 1 | 1 | 1 | 5 |
| 2 | 31 | 1 | 1 | 1 | 1 | 1 | 1 | 5 |
| 3 | 31 | 1 | 1 | 1 | 1 | 1 | 1 | 5 |
| 4 | 31 | 1 | 1 | 1 | 1 | 1 | 1 | 5 |
| 5 | 31 | 1 | 1 | 1 | 1 | 1 | 1 | 5 |
| 6 | 31 | 1 | 1 | 1 | 1 | 1 | 1 | 5 |
| 7 | 31 | 1 | 1 | 1 | 1 | 1 | 1 | 5 |
| 8 | 31 | 1 | 1 | 1 | 1 | 1 | 1 | 5 |
| 9 | 31 | 1 | 1 | 1 | 1 | 1 | 1 | 5 |
| 10 | 31 | 1 | 1 | 1 | 1 | 1 | 1 | 5 |
| 11 | 31 | 1 | 1 | 1 | 1 | 1 | 1 | 5 |
| 12 | 31 | 1 | 1 | 1 | 1 | 1 | 1 | 5 |
| 13 | 31 | 1 | 1 | 1 | 1 | 1 | 1 | 5 |
| 14 | 31 | 1 | 1 | 1 | 1 | 1 | 1 | 5 |
| 15 | 31 | 1 | 1 | 1 | 1 | 1 | 1 | 5 |
| 16 | 31 | 1 | 1 | 1 | 1 | 1 | 1 | 5 |
| 17 | 31 | 1 | 1 | 1 | 1 | 1 | 1 | 5 |
| 18 | 31 | 1 | 1 | 1 | 1 | 1 | 1 | 5 |
| 19 | 31 | 1 | 1 | 1 | 1 | 1 | 1 | 5 |
| 20 | 31 | 1 | 1 | 1 | 1 | 1 | 1 | 5 |

| **Table S6 The composition of OS medium** | |
| --- | --- |
| Abbreviation | Compound |
| ca2 | Ca2+ |
| cl | Cl- |
| cobalt2 | Co2+ |
| cu2 | Cu2+ |
| fe2 | Fe2+ |
| fe3 | Fe3+ |
| glc__D | D-Glucose |
| h2o | H2O |
| h | H+ |
| k | K+ |
| mg2 | Mg |
| mn2 | Mn2+ |
| mobd | Molybdate |
| na1 | Na+ |
| nh4 | Ammonium |
| ni2 | Ni2+ |
| o2 | O2 |
| pi | Phosphate |
| so4 | Sulfate |
| zn2 | Zn2+ |
| pi | Phosphate |
| h2 | Hydrogen |
| cobalt2 | Co2+ |
| no3 | Nitrate |
| ni2 | Nickel |

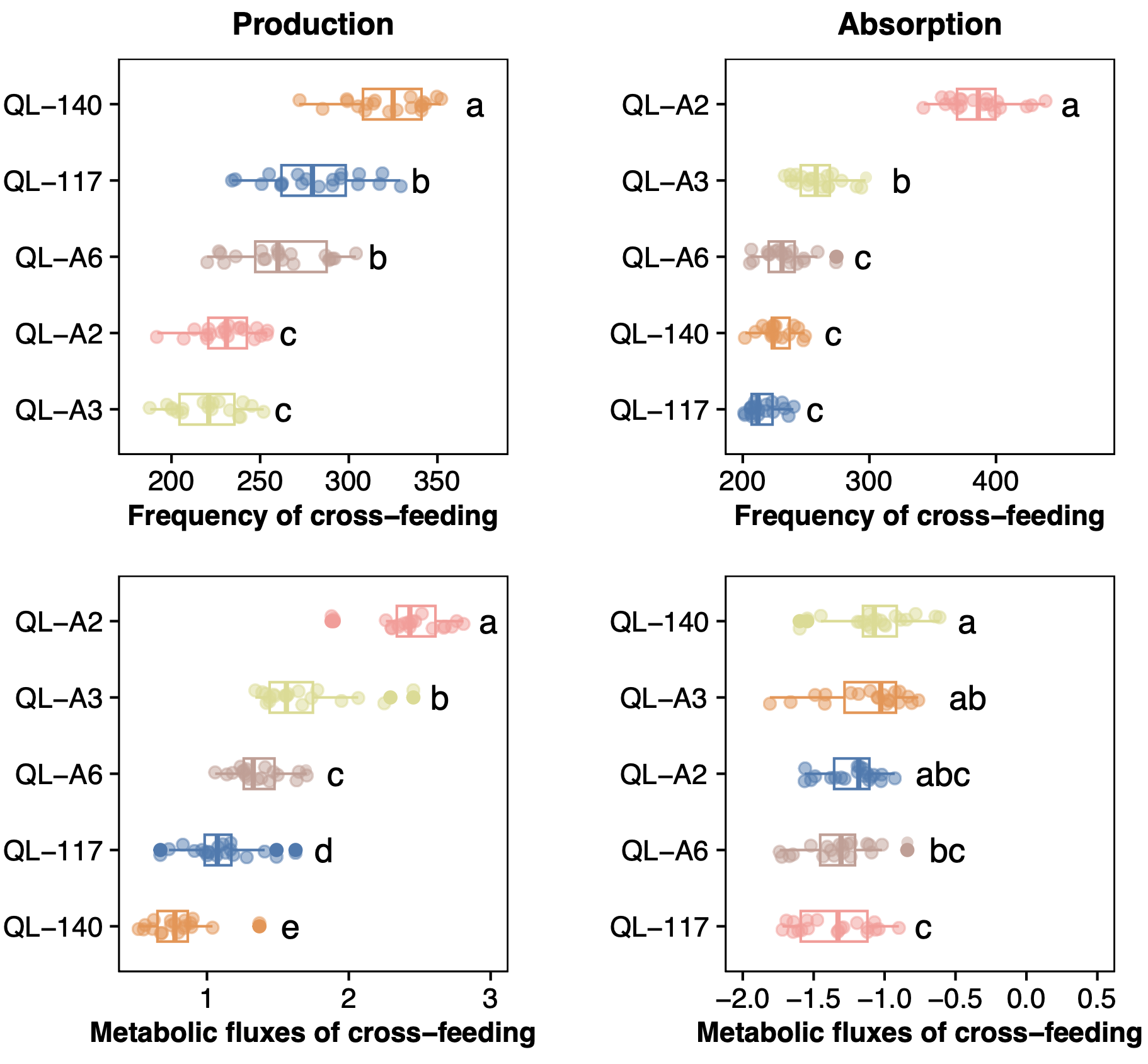

**Fig. S1** **The frequency of producing and absorbing cross-feeding metabolites within the resident community based on *in silico* simulation.**

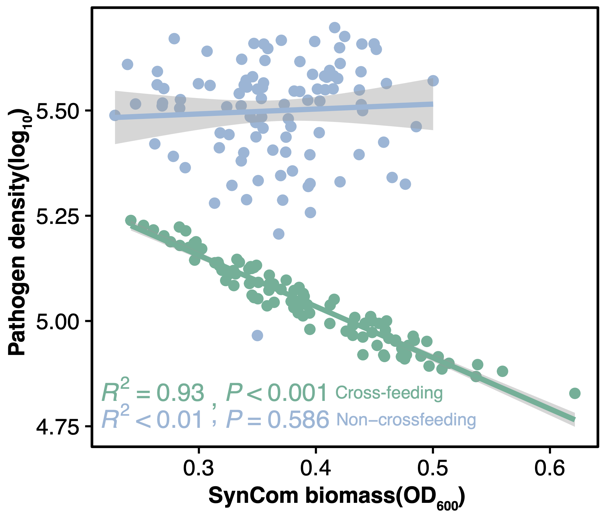

**Fig. S2** **Linear relationship between SynCom biomass and pathogen density under cross-feeding and non-crossfeeding metabolites.**
